# Tetrahydrocurcumin Suppresses Bladder Carcinogenesis via Reprogramming O-GlcNAcylation-Phosphorylation Crosstalk

**DOI:** 10.64898/2026.04.12.717994

**Authors:** Mengni Yang, Rui Li, Mengting Zhou, Yu Dong, Junning Zhao, Ruirong Tan

## Abstract

Bladder cancer exhibits high recurrence rates and limited therapeutic options in advanced stages. Although post-translational modifications (PTMs) are critical regulators of tumor biology, their systems-level remodeling during bladder carcinogenesis remains insufficiently defined. Using a BBN-induced murine bladder cancer model combined with tumor-derived organoids and bladder cancer cell lines, we observed a consistent global elevation of protein O-GlcNAcylation across tumor-associated contexts. Site-resolved quantitative proteomics revealed that this increase reflected selective redistribution rather than uniform accumulation, with differential O-GlcNAc sites nearly evenly divided between up-and down-regulated events. Integrated analyses demonstrated preferential enrichment of up-regulated O-GlcNAcylation within cytoskeleton-adhesion modules and Notch-related domains, whereas down-regulated events were more closely associated with structural maintenance and homeostatic programs. Approximately half of differentially O-GlcNAcylated proteins exhibited concurrent phosphorylation changes, indicating coordinated multi-layer PTM remodeling. Pharmacological inhibition of O-GlcNAc transferase suppressed bladder cancer cell proliferation and organoid growth, supporting a functional association between elevated O-GlcNAcylation and tumor growth. Tetrahydrocurcumin (THC) reduced global O-GlcNAcylation and induced directional remodeling of BBN-associated O-GlcNAc patterns. Cross-comparative analyses identified subsets of sites exhibiting opposite regulation trends following THC intervention. Collectively, these findings define a spatially and functionally organized PTM remodeling landscape in bladder cancer and suggest that THC exerts anti-tumor effects in part through coordinated reprogramming of O-GlcNAcylation and phosphorylation networks.

## 1 Introduction

Bladder cancer is one of the most common malignancies of the urinary system and ranks among the leading cancers in men, with a steadily increasing global disease burden. According to Global Cancer Statistics 2022 (GLOBOCAN 2022), approximately 613,791 new cases and 220,349 deaths from bladder cancer were reported worldwide in 2022, accounting for 3.1% of all newly diagnosed cancers and ranking sixth in incidence among male malignancies(1). These data highlight bladder cancer as a persistent and significant global public health challenge.

Clinically, bladder cancer is classified into non-muscle-invasive bladder cancer (NMIBC) and muscle-invasive bladder cancer (MIBC) based on muscular invasion. These two subtypes differ substantially in their mutational profiles, molecular features, and biological behaviors(2, 3). Approximately 75% of newly diagnosed cases are NMIBC, with tumors confined to the mucosa or lamina propria, whereas about 25% present with MIBC at diagnosis, which is generally associated with higher aggressiveness and poor prognosis(4, 5). Despite advances in high-throughput sequencing and multi-omics technologies, which have improved understanding of bladder cancer heterogeneity and informed precision therapeutic strategies, clinical outcomes remain unsatisfactory. Standard management of NMIBC includes transurethral resection followed by risk-adapted intravesical therapy, such as BCG immunotherapy or intravesical chemotherapy, yet 50%-70% of patients relapse within 1-3 years, with some progressing to MIBC. For MIBC, even with multimodal treatments centered on radical surgery combined with chemotherapy, radiotherapy, and more recently introduced targeted therapies and immunotherapies, the 5-year overall survival rate remains below 50%(6, 7). Comparable therapeutic limitations have also been reported in other genitourinary malignancies, including prostate cancer(8, 9), highlighting a broader challenge in achieving durable disease control through conventional oncogene-centered treatment strategies. In parallel, genetic association and replication studies have further characterized inherited susceptibility to prostate cancer in specific populations(10), yet these genomic insights have not substantially altered therapeutic outcomes.

During tumor initiation and progression, in addition to genetic alterations and transcriptional regulation, fine-tuned regulation at the protein level is equally critical. Post-translational modifications (PTMs) dynamically regulate protein activity, stability, subcellular localization, and protein-protein interactions, thereby playing essential roles in tumor-associated signaling transduction, metabolic reprogramming, and stress adaptation(11, 12). However, compared with extensive investigations at the genomic and transcriptomic levels, the global regulatory landscape and functional significance of PTMs in bladder cancer remain insufficiently characterized.

Among the diverse forms of PTMs, O-GlcNAcylation and phosphorylation represent two highly dynamic and functionally interconnected modifications that predominantly occur on serine and threonine residues. These modifications can competitively or cooperatively regulate the same or neighboring sites, thereby fine-tuning protein function and downstream signaling pathway activity(13). Accumulating evidence indicates that dysregulation of the O-GlcNAcylation-phosphorylation interplay is closely associated with multiple tumor-related processes(14); nevertheless, the overall features and biological implications of this crosstalk in bladder cancer have yet to be systematically explored. In recent years, multiple studies have demonstrated that aberrant elevation of protein O-GlcNAcylation is a prominent molecular feature of bladder cancer(15), as well as various solid tumors including colorectal, prostate, breast, and hepatocellular carcinomas(16–23). O-GlcNAcylation is a unique form of PTM in which a single N-acetylglucosamine (GlcNAc) moiety is covalently attached to the hydroxyl group of serine or threonine residues via an O-glycosidic linkage(24). This modification depends on the intracellular hexosamine biosynthetic pathway (HBP), through which approximately 2%-5% of glucose is diverted to generate the glycosyl donor UDP-GlcNAc; in addition, glutamine and glucosamine also contribute to UDP-GlcNAc biosynthesis. The addition of the GlcNAc moiety is catalyzed by O-GlcNAc transferase (OGT), whereas its removal is mediated by O-GlcNAcase (OGA). O-GlcNAcylation predominantly occurs in the nucleus, cytoplasm, and mitochondria, and more than 4,000 O-GlcNAc-modified proteins have been identified to date, encompassing a wide range of functional categories including transcription factors, kinases, histones, and cytoskeletal proteins. Owing to its rapid and reversible nature, as well as its ability to compete or cooperate with phosphorylation at specific sites, O-GlcNAcylation is extensively involved in signal transduction, gene expression regulation, cellular metabolism, proteostasis, and cell cycle control(25–29). In multiple tumor types, including bladder cancer, increased HBP flux, elevated UDP-GlcNAc levels, and enhanced global protein O-GlcNAcylation have been consistently reported, suggesting that the O-GlcNAcylation-phosphorylation interplay may constitute a metabolically sensitive and potentially targetable regulatory layer underlying tumor-associated signaling abnormalities. In line with this emerging paradigm, recent cancer studies have emphasized the therapeutic value of targeting protein regulatory networks—such as inducible cancer vulnerabilities, protein stability control, and targeted degradation(30–32)—rather than relying solely on direct inhibition of individual oncogenic enzymes or pathways. These findings underscore post-translational regulation and proteostasis as critical, actionable layers in tumor biology.

In this context, small-molecule compounds capable of modulating such dynamic PTM networks without relying on single-target inhibition may provide new opportunities to dissect and therapeutically intervene in PTM dysregulation in bladder cancer. Tetrahydrocurcumin (THC), the major active metabolite of curcumin, exhibits improved stability and bioavailability compared with its parent compound(33, 34). Accumulating evidence indicates that THC exerts anti-tumor effects through pleiotropic, multi-pathway regulatory mechanisms rather than single-target inhibition(35). For example, our previous studies showed that THC suppresses breast cancer cell proliferation and metastasis via modulation of the CYP1A/NF-κB signaling axis and remodeling of the tumor immune microenvironment(36). Consistently, THC has also been reported to attenuate colorectal cancer progression by regulating the SPP1/CD44 axis and inhibiting M2 polarization of tumor-associated macrophages(37). Meanwhile, earlier investigations have also shown that THC exerts pronounced antihyperglycemic effects in spontaneous type 2 diabetic db/db mouse models(38), further supporting its in vivo applicability and capacity for systemic regulatory modulation.

As a natural compound with pleiotropic bioactivities, THC is thought to exert its anti-tumor effects not through a single canonical target or signaling pathway, but rather through coordinated modulation of multiple regulatory nodes. Notably, our preliminary studies revealed that THC treatment reduces global protein O-GlcNAcylation levels in bladder cancer cells, suggesting a potential role for THC in modulating O-GlcNAc-related regulatory processes. Given the high metabolic sensitivity of O-GlcNAcylation and its dynamic interplay with phosphorylation, it is plausible that THC influences bladder cancer-associated signaling pathways by altering the O-GlcNAcylation-phosphorylation crosstalk. Based on this background, the present study employed a BBN-induced mouse bladder cancer model in combination with O-GlcNAcylation-phosphorylation-focused PTM proteomic profiling and in vitro cellular assays to systematically investigate the regulatory effects of THC on O-GlcNAcylation-phosphorylation modification patterns in bladder cancer, thereby providing mechanistic insights into the potential anti-tumor actions of THC in this disease context.

## 2 Materials and methods

### 2.1 Regents and instruments

BBN (N-butyl-N-(4-hydroxybutyl) nitrosamine) was purchased from Tokyo Chemical Industry (Tokyo, Japan). Tetrahydrocurcumin (THC) was obtained from Macklin (Shanghai, China). OSMI-1 and PUGNAc were purchased from MedChemExpress (MCE, Monmouth Junction, NJ, USA). The CCK-8 assay kit was supplied by Abbkine (Wuhan, China), while crystal violet stain and the BCA protein assay kit were obtained from YEASEN (Shanghai, China). Primary antibodies used in this study included anti-YAP1 and anti-β-actin (Cell Signaling Technology, Danvers, MA, USA) and an anti-O-GlcNAc mouse antibody (Sigma-Aldrich, St. Louis, MO, USA). Secondary antibodies were purchased from Cell Signaling Technology and Multi Sciences (Hangzhou, China).

### 2.2 BBN-induced bladder cancer model

All animal experiments were conducted in strict accordance with institutional guidelines and were approved by the Laboratory Animal Ethics Committee of Sichuan Academy of Chinese Medicine Sciences (Approval No. R20220303-1). Male C57BL/6 mice (4-5 weeks old, 18-20 g) were purchased from Gempharmatech and housed in a barrier facility under controlled conditions (12 h light/dark cycle, 50 ± 10% relative humidity, and 23 ± 2°C). The total experimental duration was 21 weeks. After a one-week acclimation period, mice were randomly divided into a normal control group (NC) and a bladder cancer model group (BBN) (n = 10 per group). According to previously reported methods(39, 40), the BBN group received drinking water containing 0.1% BBN starting from week 2 for 10 consecutive weeks, followed by regular drinking water for an additional 10 weeks to establish an early-to mid-stage muscle-invasive bladder cancer mouse model. During the induction period, BBN-containing drinking water was replaced twice weekly, and drinking bottles were protected from light throughout.

### 2.3 Hematoxylin and eosin (H&E)

Tumor specimens were fixed in 4% paraformaldehyde, paraffin-embedded, and cut into sections for hematoxylin and eosin (H&E) staining. Slides were captured on a Leica upright bright-field microscope coupled to the LIOO imaging system. Histological assessments were conducted by experienced clinical pathologists who were blinded to group allocation. Bladder sections were graded semi-quantitatively(41) across four pathological features—cellular degeneration, necrosis, inflammatory infiltration, and urothelial dysplasia—on a 0-4 scale, where higher scores denote greater severity. This rubric was adapted from established histopathological scoring approaches commonly applied in BBN-induced bladder cancer models and related reports(42–44).

### 2.4 Organoids construction and cultivation

Bladder tissues collected from mice with BBN-induced bladder cancer and from healthy control mice were used to generate mouse bladder tumor organoids (BTOs) and normal bladder organoids (NBOs), respectively. The organoid culture platform was established and optimized in our laboratory, with detailed procedures described in the master’s thesis of Li Jiao(45) entitled “*Construction of bladder cancer organoids for drug evaluation and tumor immune microenvironment research*”. Briefly, bladder tissues were minced and enzymatically dissociated in a digestion system containing collagenase and Y-27632 to obtain a single-cell suspension. The isolated cells were then combined with Matrigel and plated as droplets (cell-to-Matrigel ratio of 1:2, approximately 2×10⁴-3×10⁴ cells per droplet), followed by the addition of organoid culture medium. Cultures were maintained at 37 °C in a humidified incubator with 5% CO₂, with medium renewal every 3-4 days and passaging performed every 7-9 days.

### 2.5 Organoid drug sensitivity

For drug-response evaluation, organoid-derived single-cell suspensions were embedded in Matrigel and plated in 96-well plates. Specifically, 3 μL of cell suspension was mixed with 6 μL of Matrigel and dispensed into each well (final density ∼450-600 cells/well). Each condition was tested in triplicate (technical replicates). After gelation, organoid culture medium supplemented with the indicated drugs was added and cultures were maintained for 48 h. On day 3, the medium was carefully aspirated and replaced with freshly prepared drug-containing medium, and organoid morphology was documented by bright-field microscopy using a Leica microscope. The same medium renewal and imaging procedure was repeated on day 6. On day 9, organoid viability was quantified using a Cell Counting Kit-8 (CCK-8; Yeasen) according to the manufacturer’s instructions.

### 2.6 Cell cuture

The human bladder-related cell lines used in this study included the bladder cancer cell lines RT4, 5637, T24, and UMUC3, as well as the normal human urothelial cell line SVHUC. In addition, the murine bladder cancer cell line MB49 was included. RT4 originates from low-grade non-muscle-invasive urothelial carcinoma, whereas 5637, T24, and UMUC3 are derived from high-grade muscle-invasive bladder cancers with distinct molecular backgrounds. MB49 is widely used for the establishment of syngeneic bladder cancer models, and SVHUC served as a normal urothelial control in in vitro experiments. For cell culture, 5637, UMUC3, T24, MB49, and SVHUC cells were maintained in RPMI 1640 medium supplemented with 10% fetal bovine serum (FBS) and 1% penicillin-streptomycin, while RT4 cells were cultured in McCoy’s 5A medium containing the same supplements. All cells were grown at 37 °C in a humidified incubator with 5% CO₂.

### 2.7 Cell cytotoxicity assay

Cell viability was assessed using a Cell Counting Kit-8 (CCK-8; Yeasen Biotechnology). Cells were seeded into 96-well plates at a density of 5 × 10³ cells per well in 100 μL of complete medium. After cell attachment, cells were treated with the indicated concentrations of THC, OSMI-1, or PUGNAc according to the experimental design. Following 24 or 48 h of treatment, 10 μL of CCK-8 reagent was added to each well and incubated for an additional 2-4 h. Absorbance was then measured at 450 nm using a TECAN microplate reader. Wells containing medium only were used as blanks for background subtraction, and the resulting absorbance values were used to evaluate relative cell viability/proliferation.

### 2.8 Colony formation assay

Bladder cancer cells were seeded into 6-well plates at a density of 1,000 cells per well and cultured for 48 h. Cells were then treated with the indicated concentrations of THC, OSMI-1, or PUGNAc according to the experimental design and maintained in culture for a total of 9 days, with medium refreshed every 3 days. At the end of the incubation period, cells were rinsed with phosphate-buffered saline (PBS), fixed with methanol, and stained with 0.1% crystal violet. Colony numbers and colony formation were subsequently quantified using ImageJ software.

### 2.9 Sample processing for LC-MS/MS analysis

Bladder tissues were collected from experimental animals and assigned to a normal control group (NC, NC_1-NC_2), a model group (BBN, BBN_1-BBN_2), and a drug-treated group (THC, THC_1-THC_2). Each NC sample was prepared by pooling bladder tissues from four mice subjected to identical treatments, whereas each BBN and THC sample consisted of pooled tissues from three mice. Biological replicates were generated using randomized pooling. Tissues were pulverized in liquid nitrogen and lysed in buffer containing 8 M urea supplemented with protease and phosphatase inhibitors, followed by sonication. After centrifugation at 4 °C to remove insoluble debris, protein concentrations were determined using a BCA assay, and equal amounts of protein were processed for downstream analyses. Proteins were precipitated with 20% trichloroacetic acid (TCA), washed with cold acetone, and resuspended in 200 mM TEAB. Trypsin digestion was performed at an enzyme-to-protein ratio of 1:50 (w/w). Peptides were subsequently reduced with dithiothreitol and alkylated with iodoacetamide. Digested peptides were dissolved in immunoprecipitation (IP) buffer, clarified by centrifugation, and incubated overnight at 4 °C with pre-equilibrated PTM-specific antibody resin (PTM954, PTMBio) to enrich modified peptides. After extensive washing, bound peptides were eluted with 0.1% trifluoroacetic acid, pooled, and vacuum-dried. The eluates were desalted using C18 ZipTips, dried again, and subjected to LC-MS/MS analysis.

### 2.10 LC-MS/MS analysis

Peptides were separated using a NanoElute UHPLC system with mobile phase A (0.1% formic acid, 2% acetonitrile in water) and mobile phase B (0.1% formic acid in acetonitrile) at a flow rate of 450 nL/min, using a linear gradient of 6-24%B (0-40 min), 24-35%B (40-52 min), 35-80%B (52-56 min), and 80%B (56-60 min). Eluted peptides were ionized by nano-electrospray and analyzed on a timsTOF Pro mass spectrometer operated in PASEF mode. MS/MS spectra were acquired over an m/z range of 100-1700 with a spray voltage of 1.6 kV, allowing charge states of 0-5 and a dynamic exclusion of 24 s. Raw data were processed using MaxQuant (v1.6.15.0) against the Mus musculus protein database with a reverse decoy and contaminant database. Trypsin/P was specified with up to two missed cleavages. Carbamidomethyl (C) was set as a fixed modification, while oxidation (M), protein N-terminal acetylation, and serine/threonine O-glycosylation were set as variable modifications. Mass tolerances were set to 20 ppm for precursor and fragment ions. Protein and PSM false discovery rates were controlled at 1%, and only proteins identified with at least one unique peptide were retained.

### 2.11 Differential analysis of modification sites

For each modification site, relative quantitative intensities from two biological replicates in each group were averaged, and the ratio between groups was calculated and defined as the fold change (FC). Specifically, in the comparison between the BBN and NC groups, FC was calculated as the mean intensity in the BBN group divided by that in the NC group (denoted as the BBN/NC ratio). Similarly, in the THC versus BBN comparison, FC was calculated as the mean intensity in the THC group divided by that in the BBN group (denoted as the THC/BBN ratio). When a modification site exhibited a measurable intensity in one group but zero intensity in the other, FC values were assigned to avoid division by zero and infinite ratios. In the BBN vs NC comparison, sites detected only in the BBN group were assigned an FC value of 1000, whereas sites detected only in the NC group were assigned an FC value of 0.001. The same assignment rules were applied to the THC vs BBN comparison.

### 2.12 Motif analysis of modification sites

Motif analysis of modification sites was performed using the MoMo tool based on the motif-x algorithm(46). For each identified modification site, peptide sequences encompassing the modified residue and its flanking amino acids (±10 residues; ±6 residues for phosphorylation sites) were extracted as input. Background sequences were generated from all potential modification sites in the corresponding species proteome, using the same flanking residue lengths (±10 for non-phosphorylation sites and ±6 for phosphorylation sites). A sequence pattern was considered a significantly enriched motif when the number of matched peptides exceeded 20 and the statistical significance threshold was met (P < 0.000001).

### 2.13 Protein domain enrichment and annotation analysis of modified proteins

Protein domain enrichment and annotation analyses were performed using the Pfam(47) database. Fisher’s exact test was applied to assess the statistical significance of protein domain enrichment among differentially modified proteins, using all identified proteins as the background. A P value < 0.05 was considered statistically significant.

### 2.14 GO functional enrichment and annotation analysis of modified proteins

GO annotation and enrichment analyses were performed using eggNOG-mapper (v2.0), which is based on the EggNOG 6.0(48) database encompassing 12,535 organisms, including 1,322 eukaryotes, 10,756 bacteria, 457 archaea, and more than 2,500 viral genomes. GO terms were extracted from the annotation results of each identified protein and classified into cellular component, molecular function, and biological process categories. Fisher’s exact test was applied to evaluate the significance of GO enrichment for differentially modified proteins, using all identified proteins as the background. GO terms with P < 0.01 were selected for subsequent visualization and analysis.

### 2.15 KEGG pathway enrichment and annotation analysis of modified proteins

KEGG pathway enrichment and annotation analyses were performed using the Kyoto Encyclopedia of Genes and Genomes (KEGG) database(49), which categorizes pathways into metabolism, genetic information processing, environmental information processing, cellular processes, human diseases, and drug development.

Identified proteins were annotated against the KEGG pathway database using BLASTP (E-value ≤ 1e−4), and the highest-scoring hit for each sequence was retained for pathway assignment. Fisher’s exact test was applied to assess the significance of KEGG pathway enrichment among differentially modified proteins, using all identified proteins as the background. Pathways with P < 0.05 were considered significantly enriched.

### 2.16 Statistical analysis

Experimental data were analyzed using the built-in tools of GraphPad Prism (v10.1.1). Differences between two groups were evaluated using an unpaired, two-sided Student’s t-test, while comparisons among three or more groups were assessed by one-way analysis of variance (ANOVA).

## 3 Results

### 3.1 Global protein O-GlcNAcylation is elevated in bladder cancer models

Mice were administered 0.1% BBN in drinking water according to the experimental scheme shown in Figure 1A. After 22 weeks, bladder tissues were collected for histopathological analysis. H&E staining revealed pronounced pathological alterations in the BBN group, including disorganization of the lamina propria with inflammatory cell infiltration, urothelial atypia characterized by increased nuclear-to-cytoplasmic ratios, focal necrosis, and areas of squamous metaplasia with papillary hyperplasia, accompanied by reactive thickening of the muscular layer. In contrast, bladder tissues from the normal control (NC) group displayed intact mucosal and muscular architecture with clear boundaries and uniform staining. Consistently, semi-quantitative pathological scoring demonstrated significantly elevated levels of cell degeneration, necrosis, tissue inflammation, and epithelial dysplasia in BBN-treated mice compared with NC controls (Figure 1B). To evaluate whether BBN treatment was associated with changes in global protein O-GlcNAcylation levels, Western blot analysis was performed on bladder tissues. The results showed that global protein O-GlcNAcylation levels were significantly increased in BBN-treated mice relative to NC controls (Figure 1C).

**Figure 1.**
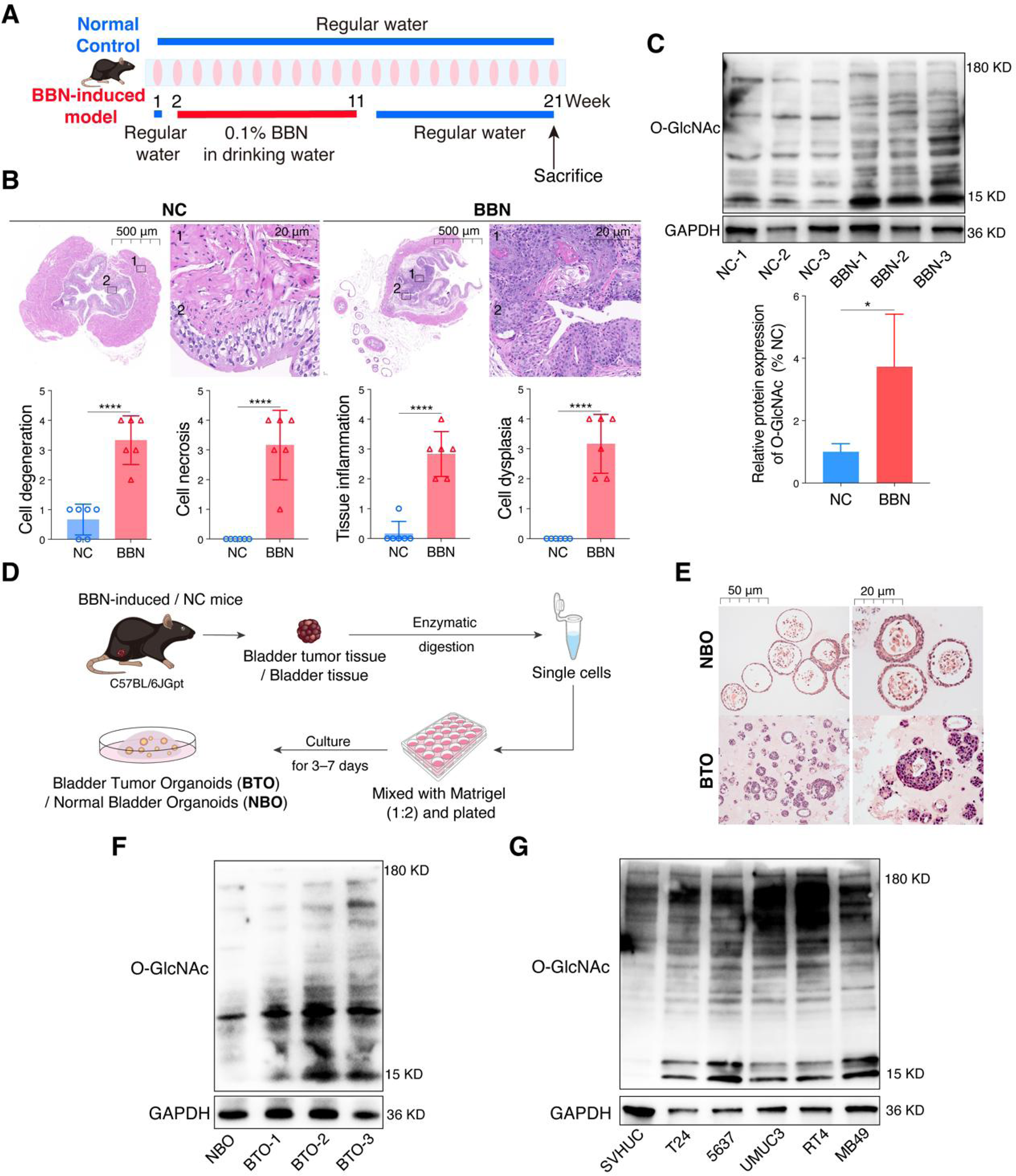
Global O-GlcNAcylation levels are increased in bladder cancer models. **(A)** Schematic of the BBN-induced bladder cancer model. **(B)** Representative H&E staining of bladder tissues from normal control (NC) and BBN-treated mice. Scale bars: 500 μm and 20 μm. Semi-quantitative analysis of pathological parameters including cell degeneration, necrosis, tissue inflammation, and dysplasia (n=6). **(C)** Western blot analysis of global O-GlcNAcylation in bladder tissues from NC and BBN mice. Quantification of relative O-GlcNAc protein levels. **(D)** Workflow for generating bladder tumor organoids (BTO) and normal bladder organoids (NBO) from mouse bladder tissues via enzymatic digestion and Matrigel-based 3D culture. **(E)** Representative histological images of NBO and BTO. Scale bars: 50 μm and 20 μm. (F) Western blot showing O-GlcNAc levels in NBO and BTOs. **(G)** Western blot analysis of global O-GlcNAcylation in human bladder epithelial cells (SV-HUC) and bladder cancer cell lines (T24, 5637, UMUC3, RT4) as well as the murine MB49 cell line. Data are presented as mean ± SD. Statistical significance was assessed using unpaired, two-tailed Student’s t-test. \**P* < 0.05, \*\**P* < 0.01, \*\*\**P* < 0.001, \*\*\*\**P* < 0.0001. * represents between indicated groups.

Subsequently, organoid models derived from BBN-induced bladder cancer tissues (BTO) and normal bladder tissues (NBO) were established (Figure 1D). H&E staining revealed that, under lower magnification (scale bar = 50 μm), NBO predominantly exhibited a regular cystic morphology with a well-defined central lumen, whereas BTO consisted mainly of densely packed cells lacking luminal structures and organized layering. Under higher magnification (scale bar = 20 μm), NBO cells displayed uniform morphology and orderly arrangement, while BTO showed pronounced cellular and nuclear atypia, including variability in nuclear size, uneven chromatin distribution, prominent nucleoli, as well as aberrant mitotic figures and multinucleated cells, resembling histopathological features commonly observed in tumor tissues (Figure 1E). In line with the observations in BBN-induced bladder tissues, Western blot analysis further demonstrated that global protein O-GlcNAcylation levels were higher in BTO than in NBO (Figure 1F). To further extend these findings, we examined global protein O-GlcNAcylation levels in a panel of bladder cancer cell lines (T24, 5637, UMUC3, RT4, and MB49) and the normal urothelial cell line SVHUC. Compared with SVHUC cells, all bladder cancer cell lines consistently exhibited elevated global protein O-GlcNAcylation levels (Figure 1G). Collectively, these results demonstrate a consistent elevation of global protein O-GlcNAcylation across multiple bladder cancer-related models, including BBN-induced bladder tissues, tumor-derived organoids, and bladder cancer cell lines, supporting a close association between enhanced O-GlcNAcylation and bladder cancer-associated pathological states.

### 3.2 Global remodeling of O-GlcNAcylation and phosphorylation landscapes in BBN-induced bladder cancer

Across multiple bladder cancer models, we first observed a consistent global elevation of protein O-GlcNAcylation. Compared with normal controls, total O-GlcNAc signals were markedly increased in BBN-induced bladder tumors as well as in additional experimental models, indicating that aberrant accumulation of O-GlcNAcylation represents a shared molecular feature of bladder carcinogenesis and is associated with tumor-associated biological contexts. However, these observations were primarily based on bulk modification levels and phenotypic readouts, and therefore did not resolve the protein-, site-, or mechanism-level organization of O-GlcNAc regulation.

To systematically characterize post-translational modification (PTM) remodeling at site resolution, we performed parallel O-GlcNAcylation and phosphoproteomic profiling of BBN-induced bladder cancer tissues (Figure 2A). Comprehensive quality control analyses demonstrated that both O-GlcNAcylated and phosphorylated peptides exhibited expected charge-state and peptide-length distributions, with highly consistent intensity profiles across samples. RSD and PCA analyses further confirmed strong reproducibility among biological replicates and clear separation between experimental groups, supporting the quantitative robustness of both datasets (Supplementary Figure 1).

**Figure 2.**
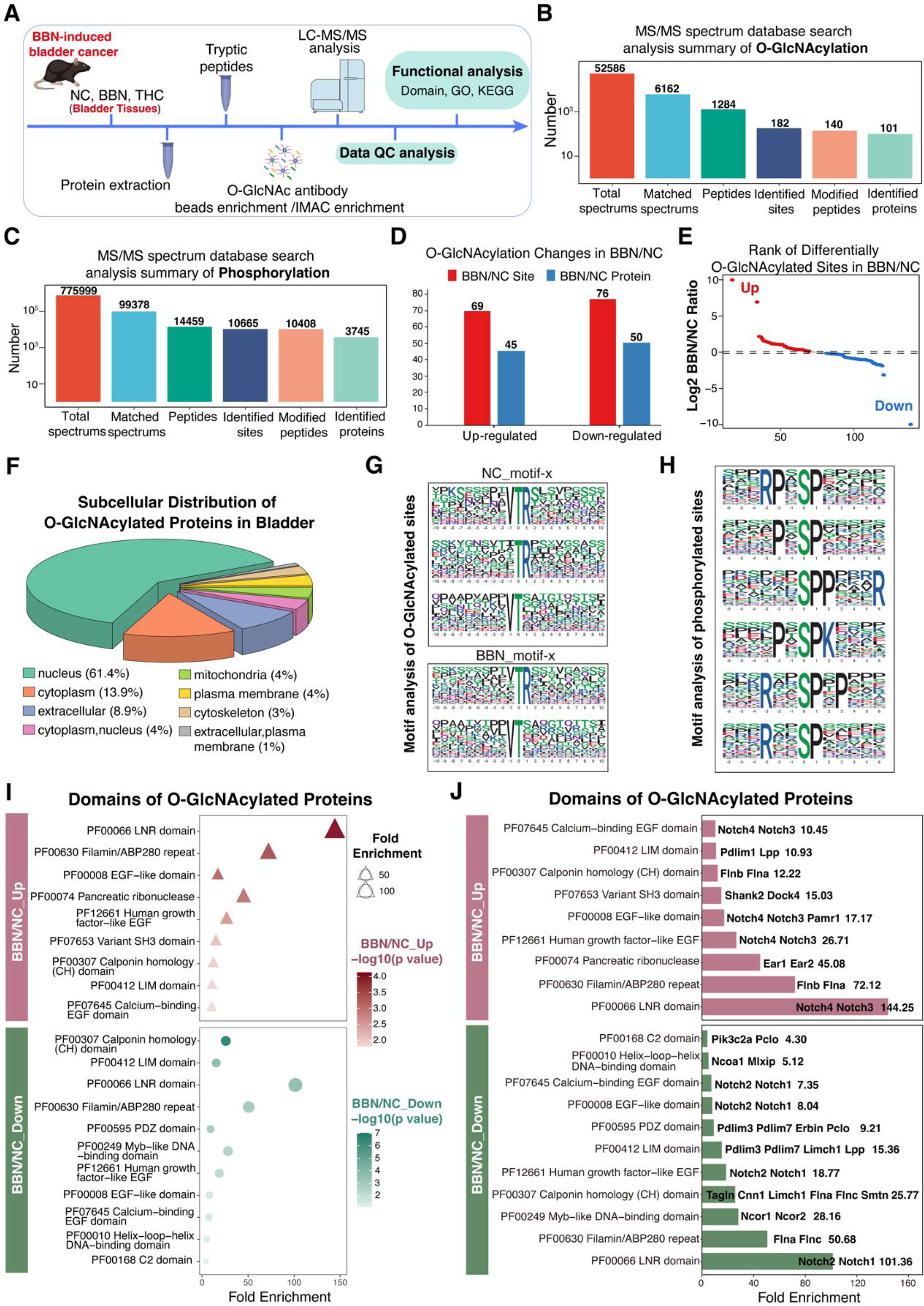
Integrated O-GlcNAcylation and phosphorylation profiling in bladder tissues. **(A)** Workflow of integrated O-GlcNAcylation and phosphorylation analysis in bladder tissues from NC, BBN, and THC-treated mice. Proteins were extracted, digested, enriched for O-GlcNAc-modified and phosphorylated peptides, and analyzed by LC-MS/MS. **(B)** Summary of MS/MS database search results for O-GlcNAcylation analysis. **(C)** Summary of MS/MS database search results for phosphorylation analysis. **(D)** Summary of differentially O-GlcNAcylated sites and proteins in the BBN *vs* NC comparisons. **(E)** Rank plots of differential O-GlcNAcylated sites in BBN *vs* NC. **(F)** Subcellular distribution of all identified O-GlcNAcylated proteins. **(G)** Motif-x-generated sequence logos of O-GlcNAcylation sites in the NC and BBN groups. **(H)** Motif-x sequence logos of phosphorylation sites. **(I)** Bubble plot showing enriched protein domains among O-GlcNAcylated proteins that were upregulated (BBN/NC_Up) or downregulated (BBN/NC_Down) in the BBN group compared with NC. **(J)** Bar plot displaying significantly enriched domains of O-GlcNAcylated proteins in the BBN/NC_Up and BBN/NC_Down groups.

Using antibody-based enrichment coupled with LC-MS/MS, we achieved comprehensive profiling of both O-GlcNAcylation and phosphorylation. In total, O-GlcNAc proteomics identified 1,284 O-GlcNAcylated peptides corresponding to 182 modification sites mapped to 101 proteins, while parallel phosphoproteomics detected 10,665 phosphorylation sites across 3,745 proteins, indicating substantial coverage depth for both PTMs (Figure 2B-C; Supplementary Data 1-2). We next examined the per-protein modification architecture to compare site burden and residue preference between the two PTMs. Overall, O-GlcNAcylation showed a low site burden, with most proteins carrying a single site (64.4%) and only a small fraction harboring ≥4 sites (5.0%) (Supplementary Figure 2A). At the protein level, O-GlcNAcylation was primarily detected on T-only proteins (67.3%), with smaller proportions in the S+T (20.8%) and S-only (11.9%) categories (Supplementary Figure 2B). Residue-stratified analyses further showed that S-site O-GlcNAcylation exhibited a relatively broader distribution of site numbers per protein, whereas T-site O-GlcNAcylation displayed a pronounced single-site predominance (Supplementary Figure 2C-D). Notably, this apparent residue distribution may be influenced by workflow-dependent enrichment and detection properties, and thus should be interpreted within the context of the applied proteomic strategy. In contrast, phosphorylation exhibited a broader multi-site profile (≥8 sites: 11.4%) (Supplementary Figure 2E) and was mainly represented by S-only (66.7%) and S+T (27.8%) proteins (Supplementary Figure 2F). Notably, T-and Y-site phosphorylation more frequently displayed high-site clustering (≥8 sites: 28.4% and 38.5%, respectively), whereas S-site phosphorylation was enriched in lower site-burden proteins (Supplementary Figure 2G-I).

Based on the aforementioned results showing elevated global O-GlcNAcylation across bladder cancer-related models, site-resolved quantification revealed that differential O-GlcNAc sites in BBN versus NC were nearly evenly split between up-and down-regulated events (Figure 2D-E, Supplementary Data 3). This pattern suggests that the increased bulk O-GlcNAc signal reflects selective redistribution and system-level remodeling of site occupancy rather than uniform accumulation across all sites. A similar bidirectional trend was observed for phosphorylation, with comparable proportions of increased and decreased sites (Supplementary Figure 3A-B, Supplementary Data 4), supporting coordinated remodeling of multiple PTM networks during tumor development.

Subcellular localization analysis showed that O-GlcNAcylated proteins were predominantly enriched in the nucleus and cytoplasm, with nuclear proteins accounting for the largest fraction (Figure 2F, Supplementary Data 5). Directional stratification revealed distinct spatial biases: up-regulated O-GlcNAcylated proteins displayed increased representation in extracellular and plasma membrane-associated compartments (Supplementary Figure 3C), whereas down-regulated proteins were further enriched in the nucleus (Supplementary Figure 3D). Phosphorylated proteins exhibited a broadly similar nuclear/cytoplasmic distribution, but with subtle compartment-specific differences between up-and down-regulated subsets, including relative enrichment of mitochondrial proteins in the down-regulated group (Supplementary Figure 3E, Supplementary Data 6). Together, these findings indicate that both O-GlcNAcylation and phosphorylation undergo spatial reorganization under BBN exposure.

Motif-x analysis revealed that O-GlcNAc sites in both NC and BBN samples preferentially occurred on serine/threonine residues and shared a conserved VTR core motif, characterized by valine at the-1 position and arginine at the +1 position. While this core preference was maintained in BBN tissues, surrounding residue composition and enrichment strength were altered, suggesting context-dependent modulation of site selection (Figure 2G, Supplementary Figure 4A-B, Supplementary Data 7). In contrast, phosphosite motifs displayed canonical proline-directed signatures centered on SP motifs with strong +1 proline preference, consistent with regulated kinase-driven phosphorylation programs (Figure 2H, Supplementary Figure 4C-D, Supplementary Data 8).

At the domain level, differentially O-GlcNAcylated proteins exhibited pronounced domain-specific enrichment. Up-regulated O-GlcNAcylation preferentially mapped to EGF-like, LIM, Filamin repeat, and calponin homology domains—modules commonly associated with cytoskeletal organization, cell-matrix adhesion, and signal integration. In contrast, down-regulated O-GlcNAcylated proteins showed distinct and weaker domain enrichment patterns (Figure 2I-J, Supplementary Data 9). Phosphorylation changes, by comparison, were biased toward signaling interaction domains such as PH, SH3, and FERM, indicating differential functional emphasis between the two PTM systems (Supplementary Figure 3F, Supplementary Data 10). Collectively, these results demonstrate that BBN-induced bladder cancer is accompanied by extensive, site-specific, and spatially organized remodeling of O-GlcNAcylation and phosphorylation networks. Rather than reflecting uniform increases in modification burden, these changes point to selective redistribution across proteins, subcellular compartments, motifs, and structural domains, providing a molecular framework for subsequent functional and mechanistic interrogation of PTM crosstalk during bladder carcinogenesis.

### 3.3 Functional enrichment analysis of O-GlcNAcylated and phosphorylated proteins in BBN vs NC

Unbiased GO enrichment analyses of proteins with increased or decreased O-GlcNAcylation in the BBN-induced bladder cancer model, performed using identical backgrounds and statistical thresholds, revealed a pronounced asymmetry in functional orientation. Proteins with increased O-GlcNAcylation preferentially converged on tumor-relevant programs, highlighting coordinated remodeling of cytoskeletal organization, cell adhesion and junctional dynamics, tissue structural behavior, and associated transcriptional regulation. These enrichments encompassed actin-myosin systems, focal adhesion-related structures, and chromatin-associated regulatory complexes, reflecting functional modules recurrently implicated in tumor progression and microenvironmental remodeling. In contrast, proteins with decreased O-GlcNAcylation were more frequently linked to developmental, physiological, or context-dependent regulatory networks, including transcriptional repression complexes and nuclear hormone receptor-associated functions, displaying a weaker direct association with tumor-promoting programs (Figure 3A, Supplementary Data 11). In parallel analyses, phosphoproteomic enrichment showed a complementary bias toward signal transduction, metabolic regulation, and stimulus-responsive pathways (Supplementary Figure 5A, Supplementary Data 12), underscoring non-redundant roles of these post-translational modification layers.

**Figure 3.**
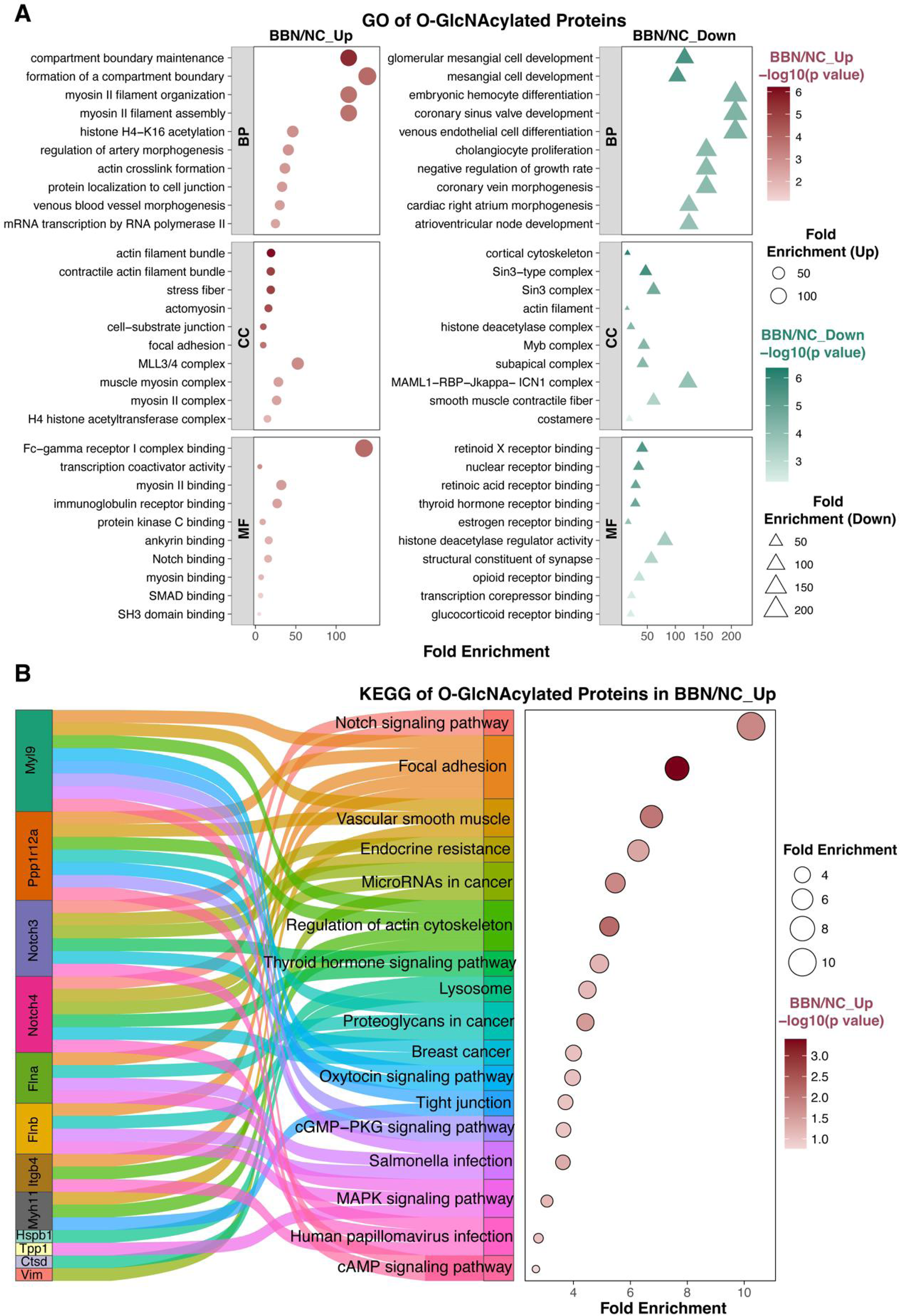
Functional enrichment analysis of differentially O-GlcNAcylated proteins in BBN *vs* NC bladder tissues. **(A)** Gene Ontology (GO) enrichment analysis of O-GlcNAcylated proteins that were significantly upregulated (BBN/NC_Up) or downregulated (BBN/NC_Down) in the BBN group compared with NC. Biological Process (BP), Cellular Component (CC), and Molecular Function (MF) categories are shown. **(B)** KEGG pathway enrichment analysis of O-GlcNAcylated proteins in the BBN/NC_Up group.

Consistent with the GO enrichment patterns, KEGG pathway analysis revealed a directional reprogramming of O-GlcNAc-associated functional networks, characterized by a clear contrast between tumor-associated remodeling and physiological homeostatic modules. Up-regulated O-GlcNAcylated proteins preferentially mapped to pathways related to adhesion-cytoskeleton organization, mechanical regulation, and Notch-associated programs, including the Notch signaling pathway, focal adhesion, regulation of the actin cytoskeleton, vascular smooth muscle contraction, and proteoglycans in cancer (Figure 3A, Supplementary Data 13). These enrichments were recurrently supported by key structural and contractile nodes such as Itgb4, Flna/Flnb, Myl9, Ppp1r12a, and Myh11, together with proteins involved in stress adaptation and proteostasis (e.g., Vim, Hspb1, Ctsd, and Tpp1). By contrast, down-regulated O-GlcNAcylated proteins were more frequently enriched in pathways associated with phosphoinositide/inositol signaling, endocrine regulation, and metabolic homeostasis, including inositol phosphate metabolism, the phosphatidylinositol signaling system, regulation of lipolysis, insulin resistance, and thyroid hormone signaling (Supplementary Figure 6). Functionally, these pathways are more closely related to physiological maintenance and metabolic balance. Notably, these enrichments largely reflect annotation-level aggregation contributed by proteins such as Notch1/Notch2 and associated transcriptional or nuclear coregulators, rather than direct pathway activation or causal regulation. In parallel, phosphoproteomic KEGG analysis showed enrichment in pathways related to metabolic adaptation and signal transduction, including calcium signaling, AMPK signaling, and hormone-or receptor-mediated pathways (Supplementary Figure 5B, Supplementary Data 14), highlighting a complementary role of phosphorylation in signaling and metabolic regulation. Although certain adhesion-and cytoskeleton-related pathways were also detected in the O-GlcNAc down-regulated set, these were primarily supported by structural maintenance components rather than canonical signaling hubs, consistent with pathway node re-wiring rather than uniform pathway-wide activation or suppression.

### 3.4 Differential O-GlcNAcylation reveals functional stratification of PPI networks in BBN-induced bladder cancer

To clarify the functional orientation of differential O-GlcNAcylation during BBN-induced bladder carcinogenesis, we performed protein-protein interaction (PPI) network analysis on proteins mapped by differential O-GlcNAc sites. PPI information was retrieved from the STRING database, with the initial network generated in STRING (Figure 4A) and subsequently visualized and optimized in Cytoscape (Figure 4B). Clear differences in network organization and functional distribution were observed between proteins associated with up-regulated and down-regulated O-GlcNAc sites.

**Figure 4.**
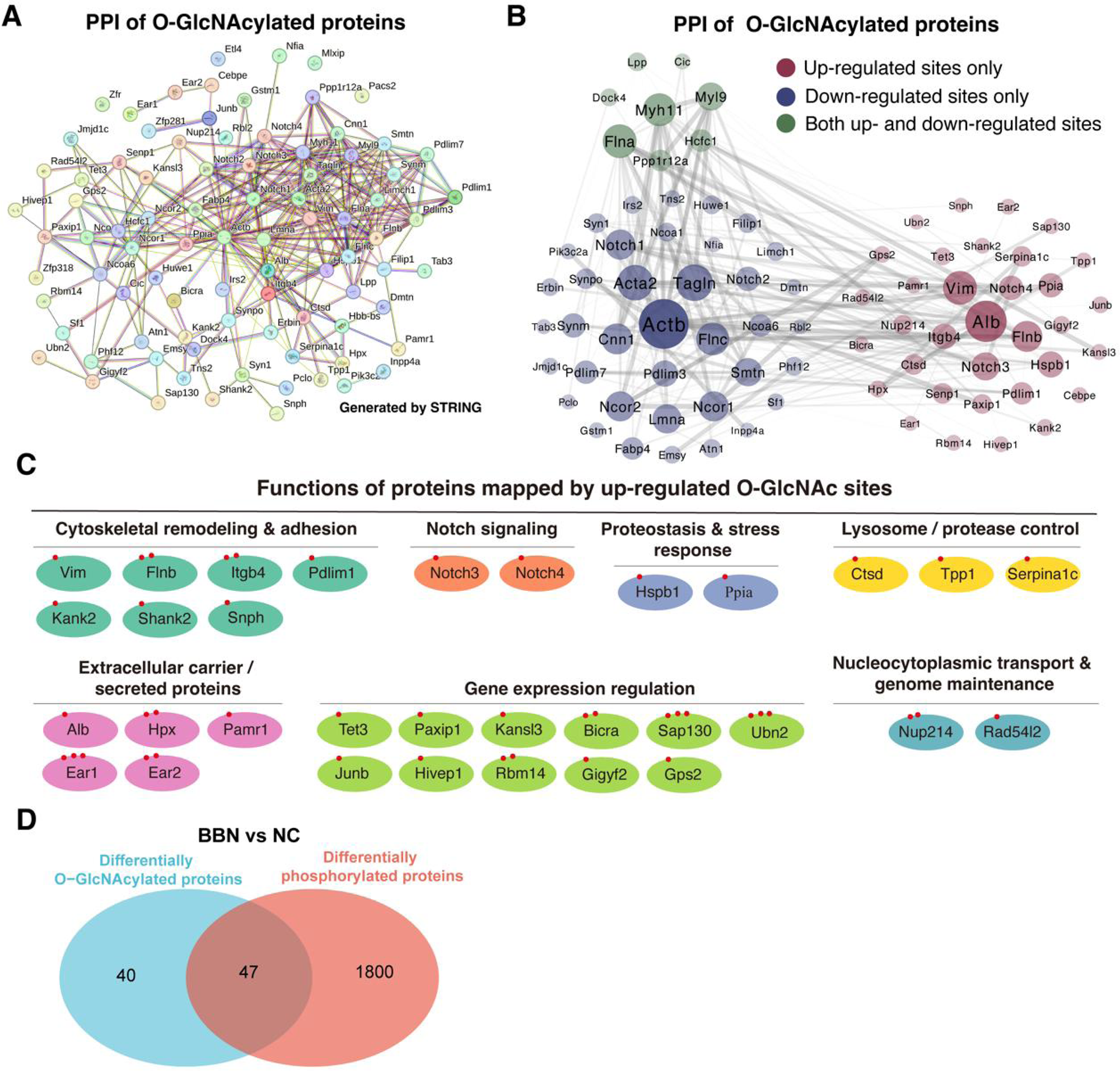
Protein-protein interaction network and functional categorization of differentially O-GlcNAcylated proteins in BBN versus NC bladder tissues. **(A)** Protein-protein interaction (PPI) network of O-GlcNAcylated proteins constructed using STRING. **(B)** Protein-protein interaction (PPI) network of O-GlcNAcylated proteins constructed using Cytoscape. Proteins harboring upregulated sites only are shown in red, those with downregulated sites only in blue, and those containing both up-and downregulated sites in green in the BBN/NC comparison. **(C)** Functional categorization of proteins associated with upregulated O-GlcNAc sites. **(D)** Venn diagram showing the overlap between differentially O-GlcNAcylated proteins and differentially phosphorylated proteins in the BBN versus NC comparison.

Proteins mapped by up-regulated O-GlcNAc sites formed densely connected modules characterized by functional annotations related to tumor-associated biological processes (Figure 4C). These modules were dominated by proteins involved in cytoskeletal remodeling and cell adhesion(50–52), including Vim, Itgb4, Flnb, and Pdlim1, which together constituted a cytoskeleton-adhesion-centered interaction subnetwork. Proteins related to cellular stress adaptation and proteostasis(53, 54) (e.g., Hspb1, Ppia, Ctsd, and Serpina1c), as well as odes annotated to phenotype-regulatory components of the Notch signaling pathway (Notch3 and Notch4)(55), were also prominently represented and showed extensive interactions with structural and regulatory proteins. In addition, several nuclear and transcription-related regulators (e.g., Tet3, Paxip1, Sap130, Ubn2, and Junb) were incorporated, extending the network to transcriptional and chromatin-associated regulation. Overall, proteins associated with up-regulated O-GlcNAc sites predominantly converged on interaction networks linked to cytoskeletal remodeling, adhesion and migration, stress adaptation, and phenotypic regulation.

In contrast, proteins mapped by down-regulated O-GlcNAc sites preferentially clustered into networks related to tissue structural maintenance and physiological homeostasis, dominated by smooth muscle contractile and cytoskeletal stability components (Acta2, Actb, Tagln, Cnn1, Smtn, and Myh11) and regulators of differentiated state and structural integrity(56–58). Notably, some proteins harbored multiple differential O-GlcNAc sites with opposite directional changes, highlighting site-specific and non-uniform regulation at the protein level. These proteins formed contractile-and support-centered interaction clusters that were clearly distinct from those associated with up-regulated O-GlcNAc sites.

We further quantified the proportion of differential O-GlcNAc-modified proteins that were accompanied by changes in phosphorylation. As shown in Figure 4D, approximately 54% (47/87) of proteins exhibiting differential O-GlcNAc modification in the BBN vs NC comparison also displayed altered phosphorylation, indicating that a substantial subset of differentially O-GlcNAcylated proteins is concurrently subject to additional layers of post-translational modification. Detailed modification site information is provided in Supplementary Data 15.

Taken together, these results demonstrate a non-random and functionally selective distribution of differential O-GlcNAc sites, with increased O-GlcNAcylation preferentially associated with proteins participating in tumor-associated interaction modules and decreased modification linked to structural maintenance and physiological homeostasis. This network-level stratification is consistent with a role for O-GlcNAcylation as an important post-translational regulatory layer accompanying tumor-associated phenotypic remodeling during BBN-induced bladder cancer development.

### 3.5 Inhibition of protein O-GlcNAcylation attenuates bladder cancer cell proliferation

Building on our earlier observations that global protein O-GlcNAcylation is markedly elevated in bladder cancer tissues, as evidenced by Western blot analysis, and that proteins mapped by up-regulated O-GlcNAcylation sites in the BBN-induced bladder cancer model are closely associated with tumor initiation and progression, we next sought to functionally assess the relationship between O-GlcNAcylation modulation and bladder cancer cell growth. To this end, bladder cancer cell lines (5637 and MB49) and bladder tumor-derived organoids (BTOs) were treated with the O-GlcNAc transferase (OGT) inhibitor OSMI-1.

CCK-8 assays revealed that OSMI-1 treatment significantly reduced the viability of both 5637 and MB49 cells in a dose-dependent manner at 24 h and 48 h, accompanied by a progressive decrease in IC50 values over time (Figure 5A-D). Consistently, colony formation assays demonstrated that pharmacological inhibition of O-GlcNAcylation markedly impaired the clonogenic capacity of bladder cancer cells compared with vehicle-treated controls (Figure 5E, F). In line with these findings in two-dimensional cultures, OSMI-1 treatment also significantly suppressed the growth of bladder tumor-derived organoids, as reflected by reduced organoid size and compromised expansion potential (Figure 5G-I).

**Figure 5.**
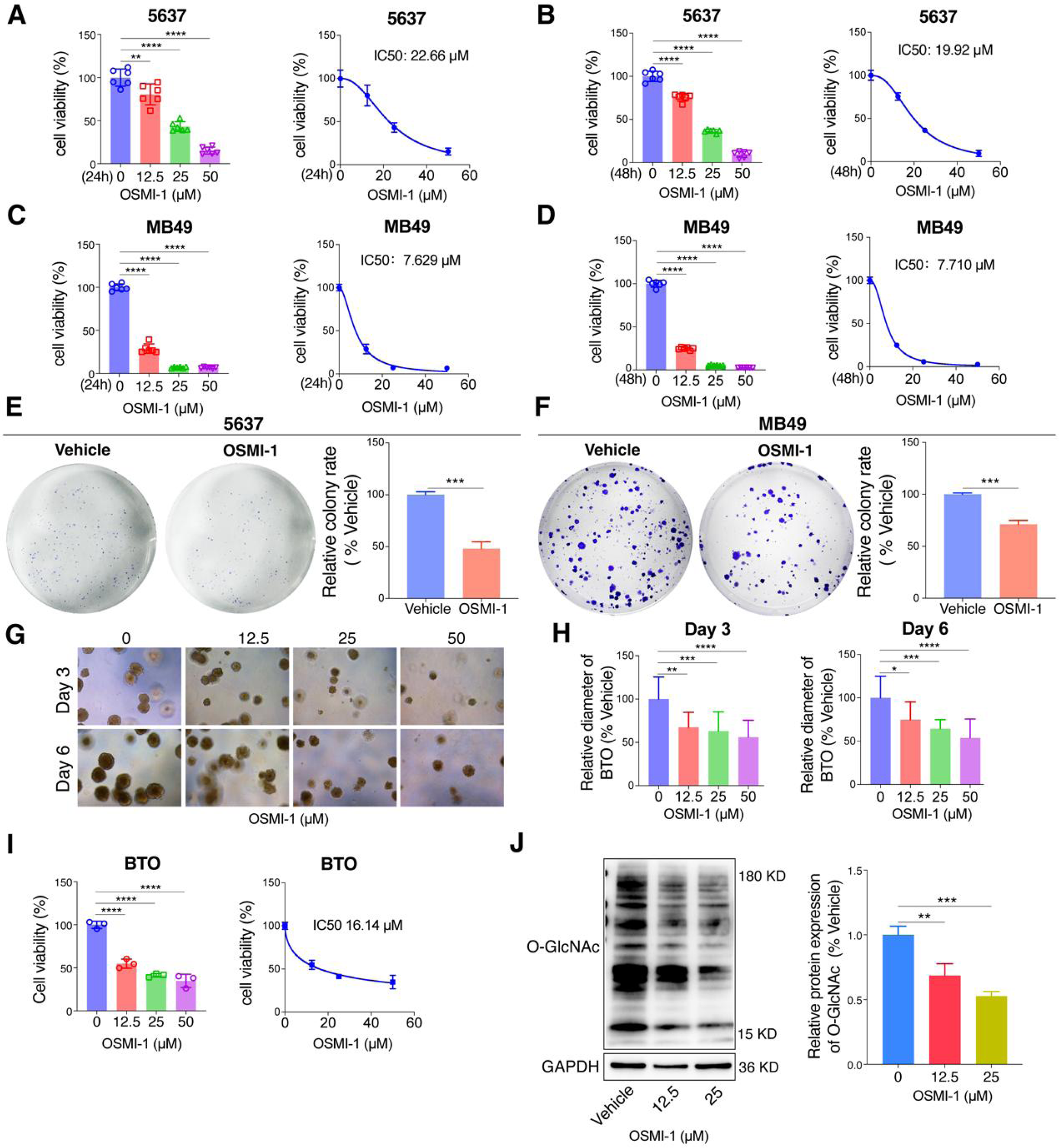
OSMI-1 suppresses bladder cancer cell growth and reduces global O-GlcNAcylation. **(A-B)** Cell viability of 5637 cells treated with OSMI-1 for 24 h (A) and 48 h (B). IC_50_ values are indicated. **(C-D)** Cell viability of MB49 cells treated with OSMI-1 for 24 h (C) and 48 h (D). IC_50_ values are indicated. **(E-F)** Colony formation assays of 5637 (E) and MB49 (F) cells treated with OSMI-1. **(G-H)** Representative images (G) and quantification (H) of bladder tumor organoids (BTOs) treated with OSMI-1 for 3 and 6 days. **(I)** Cell viability and IC_50_ analysis of BTOs treated with OSMI-1. **(J)** Western blot analysis of global O-GlcNAc levels following OSMI-1 treatment. Data are presented as mean ± SD. Statistical significance was assessed using unpaired, two-tailed Student’s t-test or one-way ANOVA followed by Tukey’s multiple comparison test. \**P* < 0.05, \*\**P* < 0.01, \*\*\**P* < 0.001, \*\*\*\**P* < 0.0001. * represents between indicated groups.

At the molecular level, Western blot analysis confirmed that OSMI-1 dose-dependently decreased global protein O-GlcNAcylation levels in bladder cancer cells, demonstrating effective suppression of intracellular O-GlcNAc modification under the applied treatment conditions (Figure 5J).

Collectively, these results demonstrate that pharmacological suppression of protein O-GlcNAcylation is consistently accompanied by impaired proliferative capacity in bladder cancer cells and tumor-derived organoids, supporting a close association between elevated O-GlcNAcylation and bladder cancer cell growth.

### 3.6 Involvement of O-GlcNAcylation in THC-mediated anti-tumor effects

Building on accumulating evidence from prior studies indicating that tetrahydrocurcumin (THC) exhibits antitumor activity(36, 37) and can modulate glycometabolic processes(38), we selected THC as the small-molecule candidate for further investigation. We subsequently evaluated its antitumor effects in bladder cancer models and examined whether these effects were accompanied by changes in O-GlcNAcylation. To define the growth-inhibitory activity of THC, CCK-8 assays were performed in 5637 and MB49 cells. THC treatment for 24 h and 48 h significantly reduced cell viability in both cell lines, and the inhibitory effect increased progressively with rising concentrations (Figure 6A-D), indicating that THC exerts a consistent antiproliferative effect in these two bladder cancer cell models.

**Figure 6.**
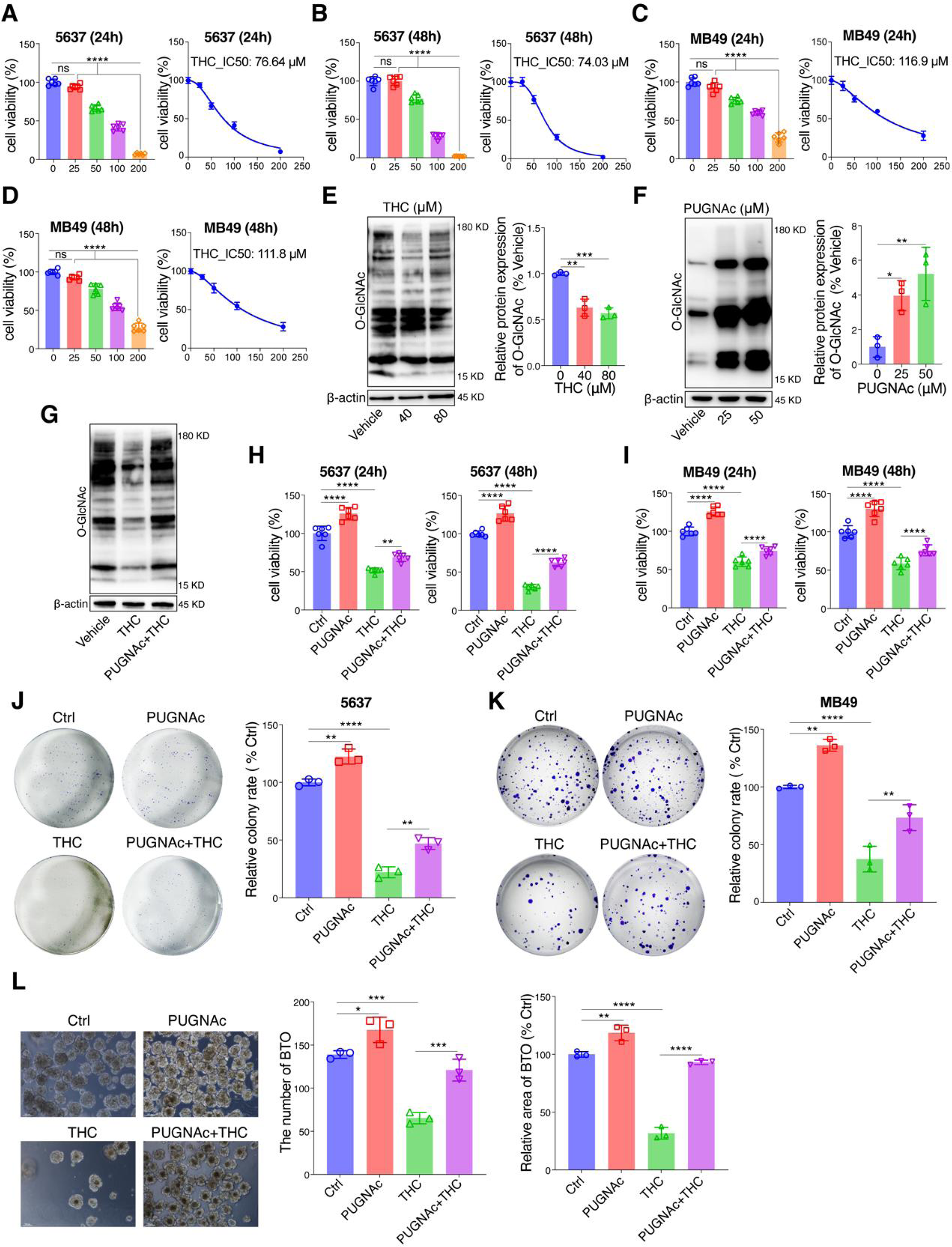
Association of O-GlcNAcylation with the anti-tumor effects of THC. **(A-B)** Cell viability of 5637 cells treated with THC for 24 h (A) and 48 h (B). IC_50_ values are indicated. **(C-D**) Cell viability of MB49 cells treated with THC for 24 h (C) and 48 h (D). IC_50_ values are indicated. **(E-F)** Western blot analysis of global O-GlcNAc levels following THC (E) or PUGNAc (F) treatment. **(G)** Western blot analysis of global O-GlcNAc levels after combined treatment with THC and PUGNAc. **(H-I)** Cell viability of 5637 (H) and MB49 (I) cells treated with THC, PUGNAc, or their combination. **(J-K)** Colony formation assays of 5637 (J) and MB49 (K) cells under the indicated treatments. **(L)** Representative images and quantification of bladder tumor organoids (BTOs) treated with THC and/or PUGNAc. Data are presented as mean ± SD. Statistical significance was assessed using one-way ANOVA followed by Tukey’s multiple comparison test. \**P* < 0.05, \*\**P* < 0.01, \*\*\**P* < 0.001, \*\*\*\**P* < 0.0001. * represents between indicated groups.

After confirming the antiproliferative phenotype, we next investigated whether this effect was associated with altered O-GlcNAc modification. Western blot analysis showed that THC markedly reduced global intracellular O-GlcNAcylation levels (Figure 6E). As a control, the OGA inhibitor PUGNAc was applied to modulate O-GlcNAcylation levels. Consistent with its reported activity(59, 60), PUGNAc treatment was associated with increased global O-GlcNAcylation (Figure 6F). Notably, under combined treatment conditions (PUGNAc + THC), the THC-induced reduction in O-GlcNAcylation was partially restored (Figure 6G), suggesting that inhibition of OGA activity can, at least in part, counteract the THC-mediated decrease in O-GlcNAcylation.

We then assessed functionally whether elevating O-GlcNAcylation would influence the growth-inhibitory effect of THC. CCK-8 assays showed that PUGNAc alone increased cell viability, whereas THC alone significantly decreased viability. Compared with THC treatment alone, combined treatment (PUGNAc + THC) partially restored cell viability in both 5637 and MB49 cells at 24 h and 48 h (Figure 6H, I). This pattern was further supported by colony formation assays: PUGNAc enhanced clonogenic capacity, whereas THC markedly suppressed colony formation. Importantly, PUGNAc partially alleviated the inhibitory effect of THC on colony formation in both cell lines under combined treatment conditions (Figure 6J, K), indicating that increased O-GlcNAcylation can attenuate, but not fully abolish, the antiproliferative effects of THC.

To more closely recapitulate tumor tissue growth, we extended these observations to bladder tumor-derived organoids (BTOs). Consistent with the cellular results, THC treatment significantly reduced organoid number and inhibited organoid growth, as reflected by a decrease in organoid area, whereas PUGNAc alone promoted organoid growth. Under combined treatment conditions, PUGNAc substantially attenuated the THC-induced suppression of organoid number and growth area, demonstrating a partial rescue effect on organoid growth (Figure 6L).

Collectively, these findings establish a clear correspondence between functional phenotypes and molecular changes: THC suppresses tumor growth in both cell-based and organoid models while concomitantly reducing global O-GlcNAcylation levels. Conversely, elevating O-GlcNAcylation through OGA inhibition partially mitigates the THC-induced downregulation of O-GlcNAcylation and, in parallel, weakens the inhibitory effects of THC on tumor growth, supporting a close functional association between O-GlcNAcylation status and THC-mediated antitumor effects rather than implying a single dominant causal mechanism.

### 3.7 Opposite remodeling of BBN-associated O-GlcNAcylation patterns following THC intervention

Given that tumor-associated upregulation of O-GlcNAcylation was observed in the BBN vs NC comparison, and that THC treatment was shown at the cellular level to reduce global O-GlcNAcylation and suppress cell growth, we next focused our analysis on the THC vs BBN comparison to examine how the O-GlcNAc landscape changes following THC intervention at the post-translational modification level. In this comparison, a total of 71 up-regulated and 69 down-regulated O-GlcNAc sites were identified, corresponding to 48 up-regulated and 49 down-regulated proteins, respectively (Figure 7A; Supplementary Data S16). Ranking analysis based on the magnitude of modification changes (Figure 7B) indicated that THC treatment was associated with a broad remodeling of the O-GlcNAcylation landscape, rather than affecting only a limited number of sites.

**Figure 7.**
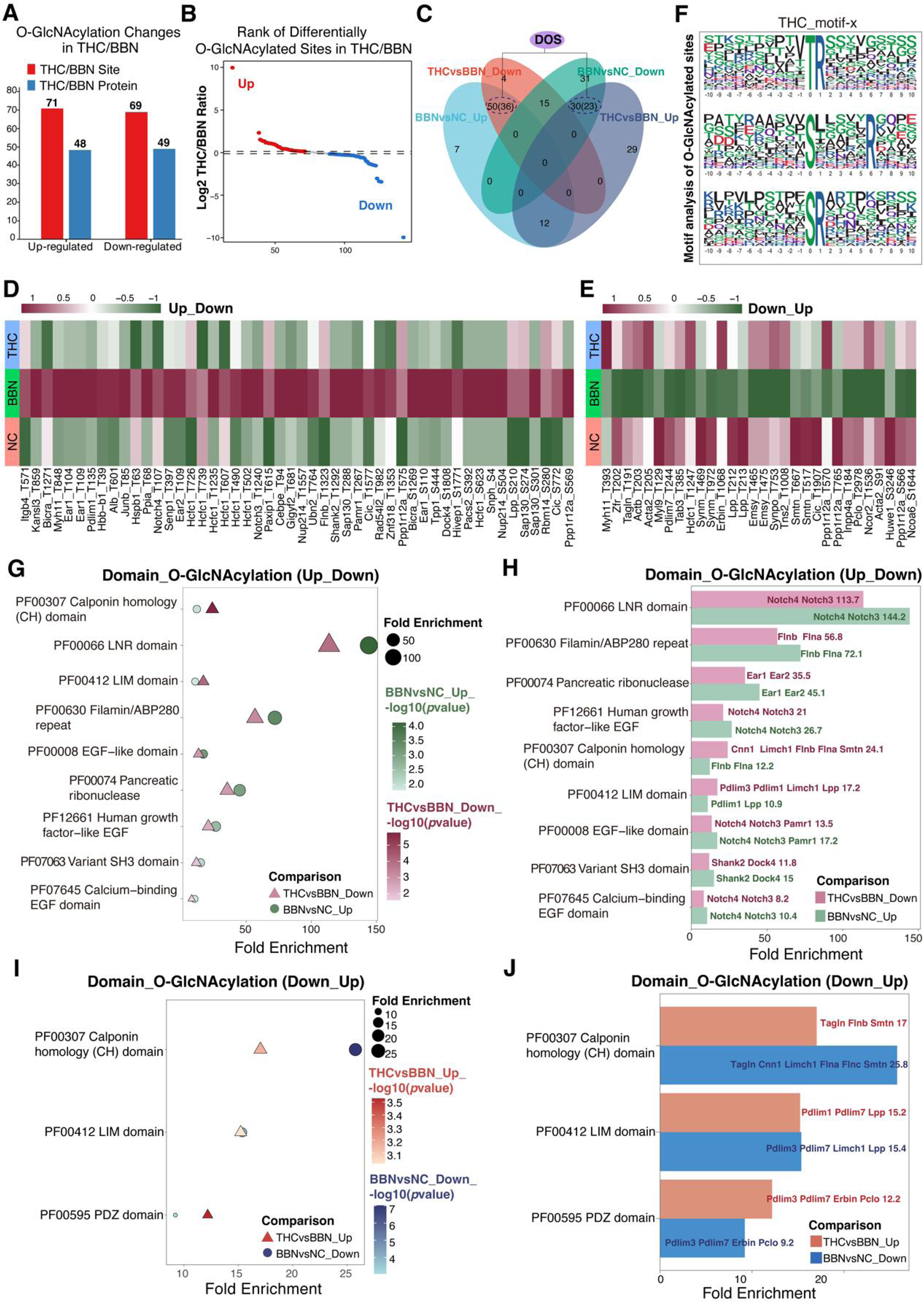
THC remodels the O-GlcNAcylation landscape in BBN-induced bladder cancer. **(A)** Summary of differentially O-GlcNAcylated sites and proteins in the THC *vs* BBN comparisons. **(B)** Rank plots of differential O-GlcNAcylation sites in THC *vs* BBN. **(C)** Intersection analysis of directionally regulated differential O-GlcNAc sites (DOS) across BBN vs NC and THC vs BBN (Up/Down patterns). **(D-E)** Heatmaps showing representative O-GlcNAcylation sites exhibiting opposite regulatory patterns across groups. Up_Down pattern: sites increased in BBN relative to NC but decreased after THC treatment (D). Down_Up pattern: sites decreased in BBN relative to NC but restored following THC treatment (E). **(G-H)** Protein domain enrichment analysis for O-GlcNAcylated proteins showing the Up_Down pattern. Bubble plot displaying enriched domains based on fold enrichment and statistical significance (G). Bar plot summarizing the enriched domains and representative proteins within each domain (H). **(I-J)** Protein domain enrichment analysis for O-GlcNAcylated proteins showing the Down_Up pattern. Bubble plot showing enriched domains associated with restored O-GlcNAcylation after THC treatment (I). Bar plot highlighting representative proteins within each enriched domain (J).

To more clearly delineate the directionality of THC-associated regulation relative to BBN-induced changes, we performed a cross-comparison based on regulation trends between the two datasets (Figure 7C). In this analysis, 50 differential O-GlcNAc sites (DOS), mapping to 36 proteins, were up-regulated in BBN vs NC but down-regulated in THC vs BBN (Up_Down; Figure 7D). Conversely, 30 DOS (corresponding to 23 proteins) were down-regulated in BBN vs NC but up-regulated following THC treatment (Down_Up; Figure 7E). Together, these sites define a subset of O-GlcNAcylation events that display opposite directional changes between carcinogen induction (BBN) and THC treatment, indicating a reversal-oriented shift of BBN-associated O-GlcNAc modification patterns under THC intervention. As a complementary reference, we also summarized the overall changes in global phosphorylation (Supplementary Figure 7A-C; Supplementary Data S17).

Subcellular localization analysis indicated that proteins with oppositely regulated O-GlcNAc modifications were mainly nuclear and cytoplasmic, with additional distribution across mitochondria, cytoskeleton, plasma membrane, and extracellular compartments. It should be noted that this distribution reflects annotation-level localization of the modified proteins rather than the subcellular sites of O-GlcNAc modification per se. The Up_Down group showed a stronger nuclear/cytoplasmic bias with more membrane/extracellular representation (Supplementary Figure 7D), whereas the Down_Up group remained nucleus-dominant but was relatively more enriched in mitochondrial and cytoskeletal compartments (Supplementary Figure 7E). This distribution pattern suggests that the reversal effects associated with THC treatment may influence not only nuclear regulatory processes but also metabolic states and cellular structural organization. A similar nuclear-cytoplasmic enrichment pattern was observed for differentially phosphorylated proteins (Supplementary Figure 7F).

Sequence context analysis using Motif-x further revealed characteristic features of THC-associated O-GlcNAc sites (Figure 7F; Supplementary Figure 7G-H). The modified residue was predominantly serine (S) at position 0, with flanking regions displaying a low-complexity sequence enriched in serine/threonine residues. Within the-6 to +6 window, small amino acids (A/G/V) were frequently observed, and a moderate enrichment of proline at upstream proximal positions (-1/-2) was also evident.

Extending the analysis from modification sites to protein-level organization, we performed domain enrichment analysis on proteins carrying differential O-GlcNAc modifications. In the Domain Up_Down group (up-regulated in BBN vs NC and down-regulated in THC vs BBN), significantly enriched domains included Notch-associated LNR domains (Notch3/Notch4), Filamin/ABP280 repeat domains (FLNA/FLNB), multiple EGF and calcium-binding EGF-like domains, as well as Variant SH3 domains (Figure 7G-H). Collectively, these domains point toward directional remodeling of Notch-related signaling modules and cytoskeleton/adhesion-associated structural frameworks. In contrast, the Domain Down_Up group (down-regulated in BBN vs NC and up-regulated following THC treatment) exhibited a more focused enrichment profile, dominated by Calponin homology (CH) domains, LIM domains, and PDZ domains (Figure 7I-J). These domains are broadly associated with cytoskeletal dynamics, adhesion complexes, and protein-protein interaction platforms, and their relative enrichment following THC treatment suggests partial recovery of structural and interaction networks that are attenuated under BBN-induced conditions, rather than complete restoration to the normal state.

Finally, to place these O-GlcNAc-associated domain patterns within a broader post-translational modification context, we conducted parallel domain enrichment analyses on differentially phosphorylated proteins (Supplementary Figure 8A-B). Overall, limited domain overlap was observed in the BBN-up/THC-down phosphorylation trend (Supplementary Figure 8C). In contrast, a broader set of shared domains was evident in the BBN-down/THC-up trend, prominently including LIM, CH, and PDZ domains, along with Lamin tail and Variant SH3 domains (Supplementary Figure 8D). These findings indicate that modules related to cytoskeletal organization, adhesion, and protein interaction more strongly characterize phosphorylation events that are reduced by BBN induction but show directional recovery tendencies following THC treatment.

### 3.8 Functional reversal of tumor-associated O-GlcNAcylation and phosphorylation signatures following THC treatment

In the preceding BBN vs NC comparison, we observed a clear functional divergence between O-GlcNAcylation and phosphorylation: proteins mapped from up-regulated modification sites were preferentially associated with tumor-related structural remodeling and oncogenic signaling programs, whereas those mapped from down-regulated sites were more closely linked to physiological structural maintenance and homeostatic regulation. Building on this framework, we next integrated GO and KEGG enrichment analyses to systematically evaluate functional and pathway features associated with differential O-GlcNAcylation and phosphorylation in the THC vs BBN comparison, with the aim of assessing whether THC treatment induces directionally opposite functional remodeling trends relative to these abnormal regulatory patterns.

At the O-GlcNAcylation level, differentially modified proteins following THC treatment exhibited functional enrichment trends that were directionally opposite to those observed during BBN induction in both the Up_Down (up-regulated in BBN and down-regulated by THC) and Down_Up (down-regulated in BBN and restored by THC) directions (Supplementary Figure 9A-B). Specifically, Up_Down-associated proteins were primarily enriched in processes related to cell adhesion, cytoskeletal remodeling, and junctional structures, with GO cellular component terms highlighting contractile and junctional elements (such as actin filament bundles, Z discs, sarcomeres, and stress fibers), while GO molecular function terms were dominated by regulatory binding activities, including receptor binding, transcription factor/cofactor binding, and protein binding (Figure 8A). In contrast, Down_Up-associated proteins were more closely linked to cytoskeletal organization, maintenance of the contractile apparatus, and structural stability, with persistent enrichment of contractile fiber-and sarcomere-related structures (Figure 8B).

**Figure 8.**
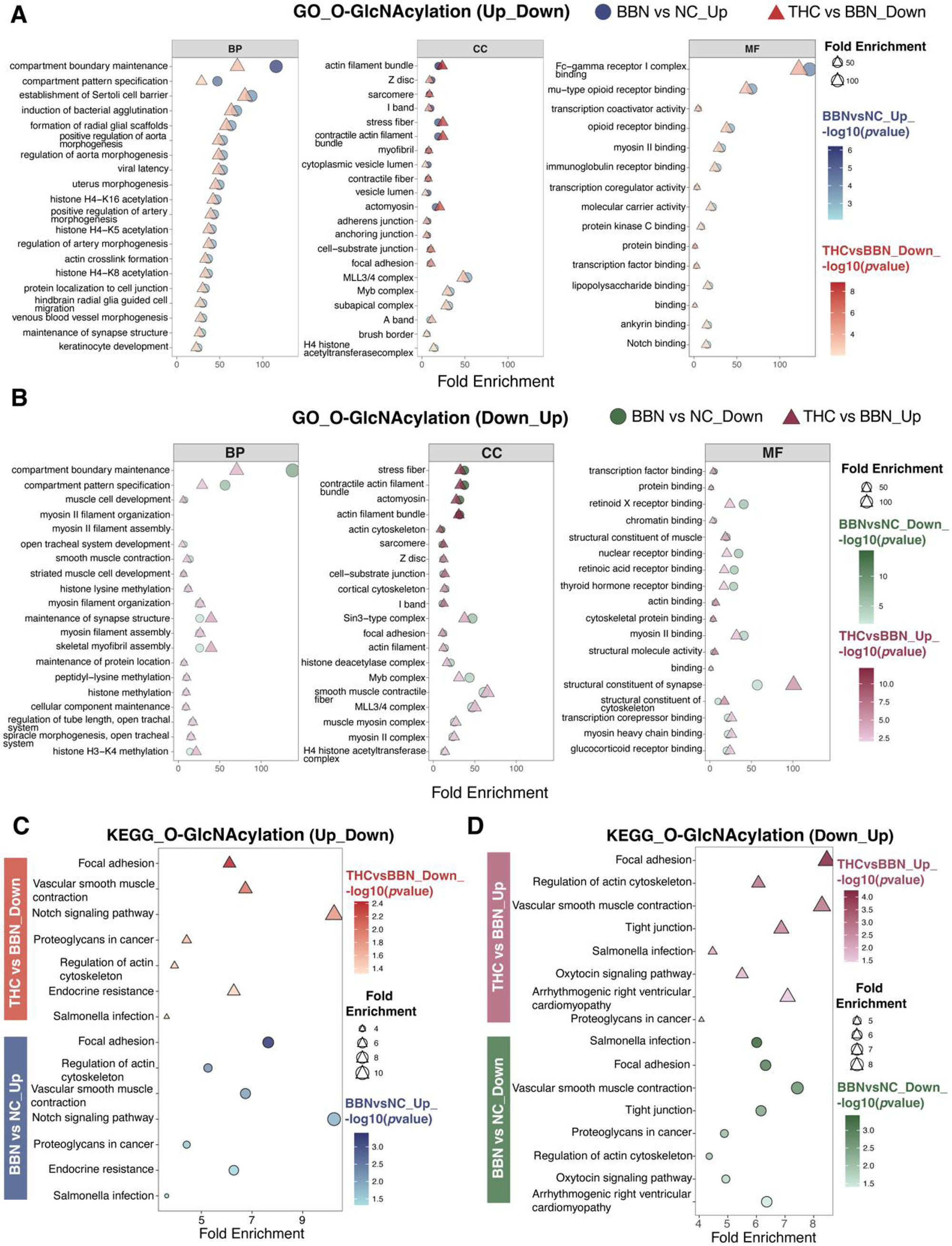
GO and KEGG enrichment analyses of differentially O-GlcNAcylated proteins. **(A-B)** GO enrichment results for the Up_Down (BBN vs NC_up and THC vs BBN_down) (A) and Down_Up (BBN vs NC_down and THC vs BBN_up) (B) patterns, shown across BP, CC, and MF categories. **(C-D)** KEGG pathway enrichment for the Up_Down (C) and Down_Up (D) patterns.

Consistent with the GO results, KEGG analysis of O-GlcNAcylated proteins revealed highly overlapping pathway sets in both the Up_Down and Down_Up modes (Supplementary Figure 9C-D). These pathways were centered on “adhesion-cytoskeleton-junction” programs, including focal adhesion, regulation of actin cytoskeleton, tight junction, and vascular smooth muscle contraction, accompanied by coordinated directional modulation of tumor-related pathways such as the Notch signaling pathway and proteoglycans in cancer (Figure 8C-D). Collectively, these findings indicate that the structural and mechanical programs aberrantly activated in the BBN model at the O-GlcNAcylation level exhibit directionally opposite enrichment trends following THC treatment, rather than complete normalization to the NC state.

By comparison, the phosphorylation layer displayed a broader and more functionally heterogeneous regulatory profile (Supplementary Figure 9E-F). GO enrichment analysis showed that Up_Down-associated terms were more strongly oriented toward immune-related processes, including regulation of T-cell proliferation, IL-2 production, and leukocyte migration, whereas Down_Up-associated terms were more closely associated with cytoskeletal organization and contractile programs (Supplementary Figure 10A-B). Further KEGG analysis indicated that differential phosphorylation also involved metabolic and energy homeostasis-related pathways as well as multiple canonical signaling axes, including MAPK, Ras, mTOR, HIF-1, and ErbB signaling pathways (Supplementary Figure 11), reflecting broader pathway-level rebalancing rather than uniform reversal across signaling networks.

Taken together, within the regulatory framework established in the BBN vs NC comparison—characterized by up-regulation of tumor-promoting modifications and down-regulation of physiological structural modifications—THC treatment exerted a consistent opposite-direction remodeling trend at both the O-GlcNAcylation and phosphorylation levels. Specifically, THC was associated with attenuation of structure-and signaling programs linked to tumor progression, while promoting relative enrichment of functional modules associated with physiological structural maintenance and homeostatic regulation. These results support the notion that THC exerts anti-bladder cancer effects by reshaping, rather than completely reversing, the O-GlcNAcylation and phosphorylation landscapes across multiple layers of post-translational modification networks.

## 4 Discussion

In this study, we systematically characterized O-GlcNAcylation remodeling in bladder cancer by integrating histopathological validation, site-resolved post-translational modification (PTM) profiling, functional perturbation, and pharmacological intervention across multiple experimental models. We demonstrate that global protein O-GlcNAcylation is consistently elevated in BBN-induced bladder tumors, tumor-derived organoids, and bladder cancer cell lines, supporting the view that enhanced O-GlcNAcylation represents a conserved molecular feature associated with bladder cancer-related pathological states. This observation is consistent with extensive evidence from other malignancies indicating that O-GlcNAcylation functions as a nutrient-and stress-responsive PTM frequently upregulated in cancer and linked to tumor cell proliferation, survival, and adaptive stress responses(61).

Despite the pronounced increase in global O-GlcNAcylation, site-resolved proteomic analysis revealed a balanced distribution of up-and down-regulated O-GlcNAc sites in BBN-induced tumors. This finding highlights that tumor-associated increases in bulk O-GlcNAc signals can arise from selective redistribution of modification events rather than uniform site-wide accumulation, a pattern also observed in other cancer and stress-adaptive contexts(62, 63). Parallel phosphoproteomic profiling further revealed coordinated but non-redundant remodeling of phosphorylation networks, underscoring the complementary roles of these PTM layers in shaping tumor-associated signaling states. Together, these results support the concept that resolving PTM regulation at site-level resolution is essential for interpreting how global modification changes translate into functional reprogramming.

Functional enrichment and interaction network analyses further indicated a clear functional stratification of O-GlcNAc-modified proteins. Proteins with increased O-GlcNAcylation preferentially clustered into modules related to cytoskeletal remodeling, cell adhesion, junctional organization, and stress-adaptive programs, whereas proteins with decreased O-GlcNAcylation were more closely linked to structural maintenance and physiological homeostasis. These findings align with growing evidence that O-GlcNAcylation can modulate cytoskeletal dynamics, mechanotransduction, and cell-matrix interactions—processes central to tumor invasion and microenvironmental remodeling(64). Notably, enrichment and network analyses reflect annotation-level aggregation and module-level organization rather than direct measurements of signaling output; therefore, they are most appropriately interpreted as indicators of functional trends and network reconfiguration rather than definitive evidence of pathway activation or suppression.

Consistent with these network-level patterns, pharmacological inhibition of O-GlcNAcylation using the OGT inhibitor OSMI-1 was accompanied by impaired proliferation and clonogenic growth in bladder cancer cells and tumor-derived organoids, supporting a close functional association between elevated O-GlcNAcylation and bladder cancer cell fitness(65–67). Building on this framework, we identified tetrahydrocurcumin (THC) as a small-molecule modulator that suppresses bladder cancer cell and organoid growth while concomitantly reducing global O-GlcNAcylation levels. Rescue experiments using the OGA inhibitor PUGNAc demonstrated that elevating O-GlcNAcylation partially attenuates the antiproliferative effects of THC, indicating that O-GlcNAcylation constitutes one contributing layer within a broader network of THC-responsive pathways. The partial rescue is consistent with the pleiotropic activities reported for curcumin derivatives and suggests that THC likely reshapes tumor-associated states through multi-target and multi-layer regulation rather than a single dominant mechanism.

At the PTM landscape level, THC treatment induced directionally opposite remodeling of O-GlcNAcylation patterns relative to BBN-induced alterations, attenuating tumor-associated structural and mechanical modules while promoting relative enrichment of components linked to physiological structural organization. Parallel phosphorylation analyses revealed broader rebalancing across immune, metabolic, and signaling pathways, suggesting that THC reshapes interconnected PTM networks in a directionally corrective manner. Together, these observations support a model in which O-GlcNAcylation functions as an important regulatory layer linking metabolic state to tumor-associated structural and signaling programs in bladder cancer, and highlight the potential therapeutic relevance of targeting PTM network remodeling rather than individual oncogenic nodes.

As our findings collectively support a PTM network-level remodeling model in bladder cancer, it is also important to interpret the current evidence within its methodological scope and to clarify the key remaining questions for future investigation. Despite the strengths of this study, several limitations should be acknowledged. First, although pharmacological modulation of O-GlcNAcylation using OSMI-1 and PUGNAc provided functional insight, genetic approaches targeting OGT or OGA would further strengthen causal inference at the molecular level. Second, enrichment and network analyses reflect annotation-level aggregation of differentially modified proteins rather than direct measurements of pathway activity, and thus should be interpreted as indicators of functional trends rather than definitive evidence of signaling activation or suppression. Third, while THC clearly modulated O-GlcNAcylation-associated phenotypes, the precise molecular targets through which THC influences O-GlcNAc cycling and PTM crosstalk remain to be elucidated. Future studies integrating genetic manipulation, biochemical target identification, and in vivo therapeutic evaluation will be important to further define the mechanistic basis and translational potential of PTM-targeted interventions in bladder cancer. Overall, by delineating selective redistribution of tumor-associated O-GlcNAcylation across protein networks and demonstrating that these signatures can be directionally counteracted by THC, this study provides a framework for understanding PTM-driven reprogramming in bladder carcinogenesis and motivates future efforts toward mechanism-guided PTM network targeting.

## Supplementary Information

### This file includes

Supplementary Figures 1 to 11

## Acknowledgments

The authors thank Hangzhou Jingjie PTM BioLab Co., Ltd. (Hangzhou, China) for technical support in O-GlcNAcylation and phosphorylation proteomics analyses.

## Funding

This research is supported by the National Natural Science Foundation of China (Grant No. 82204695).

## Competing interests

The authors declare that they have no competing interests.

## Author contributions

**Mengni Yang:** Data curation, Formal analysis, Investigation, Methodology, Software; Visualization, Writing - original draft. **Rui Li:** Data curation,Visualization, Formal analysis. **Mengting Zhou:** Formal analysis, Validation. **Yu Dong:** Validation, Project administration. **Junning Zhao:** Conceptualization, Methodology, Supervision. **Ruirong Tan:** Conceptualization, Funding acquisition, Writing - review & editing.

## Ethics approval

All animal experiments were conducted in accordance with institutional guidelines and approved by the Laboratory Animal Ethics Committee of Sichuan Academy of Chinese Medicine Sciences (Approval No. R20220303-1).

## Consent for publication

Not applicable.

---

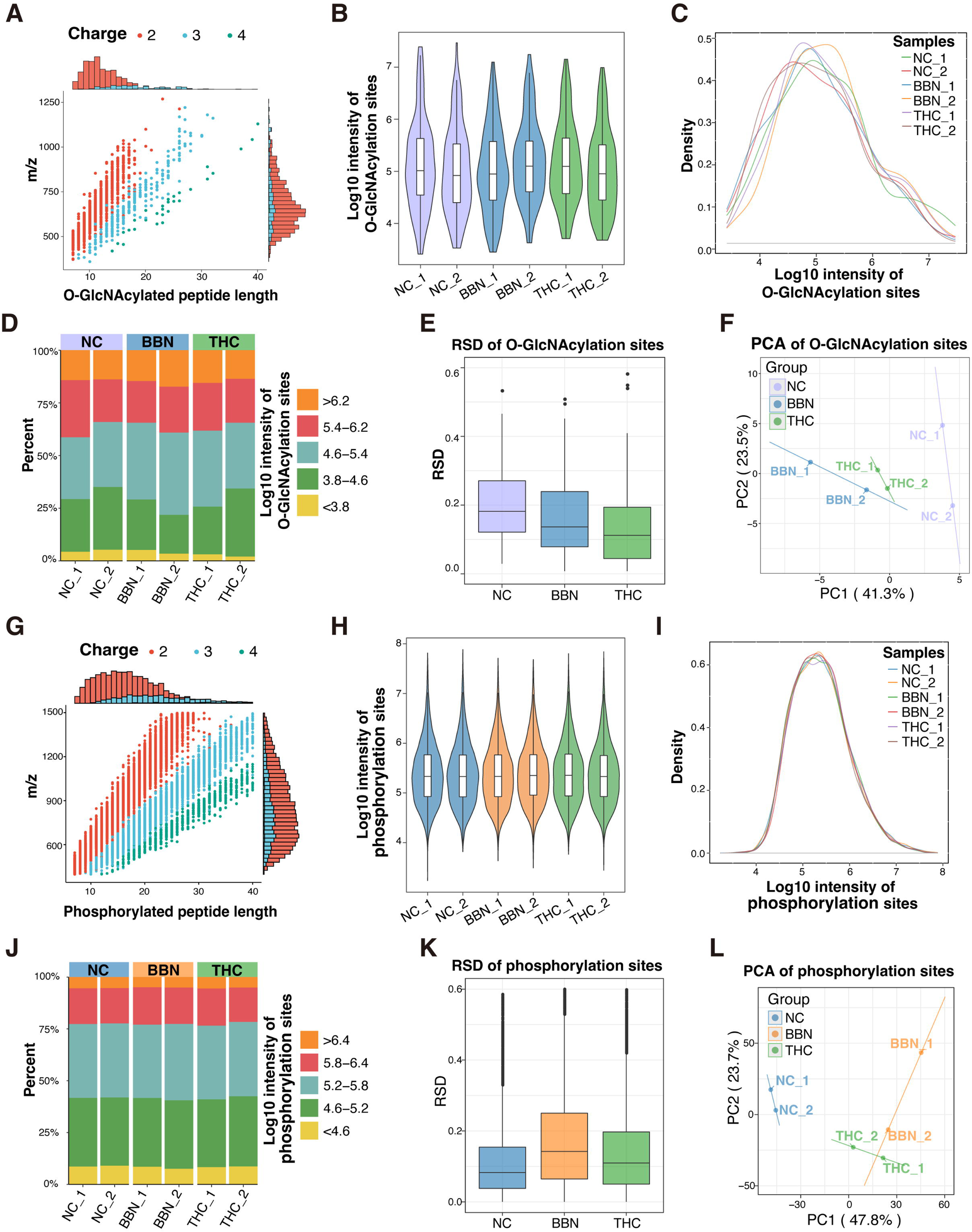

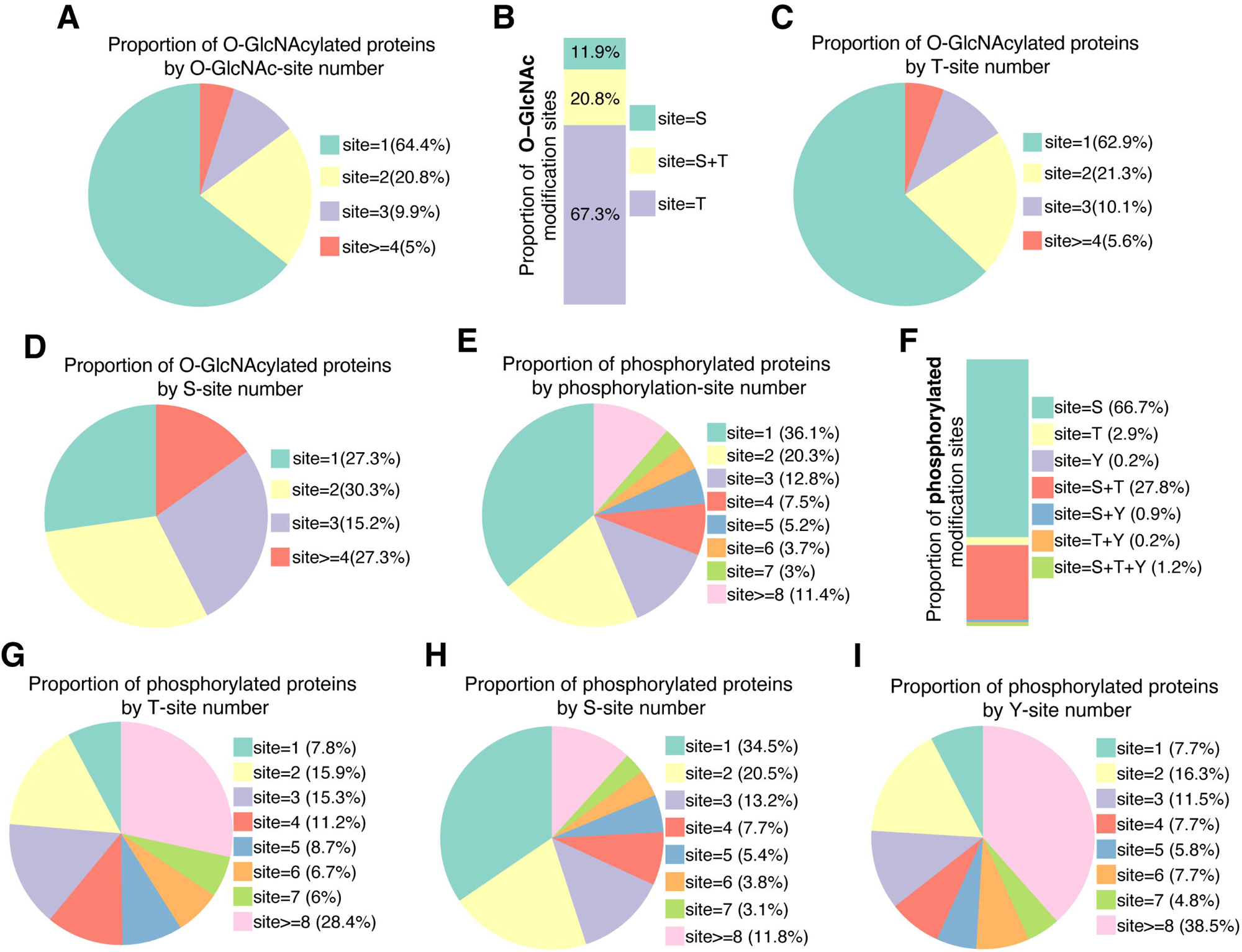

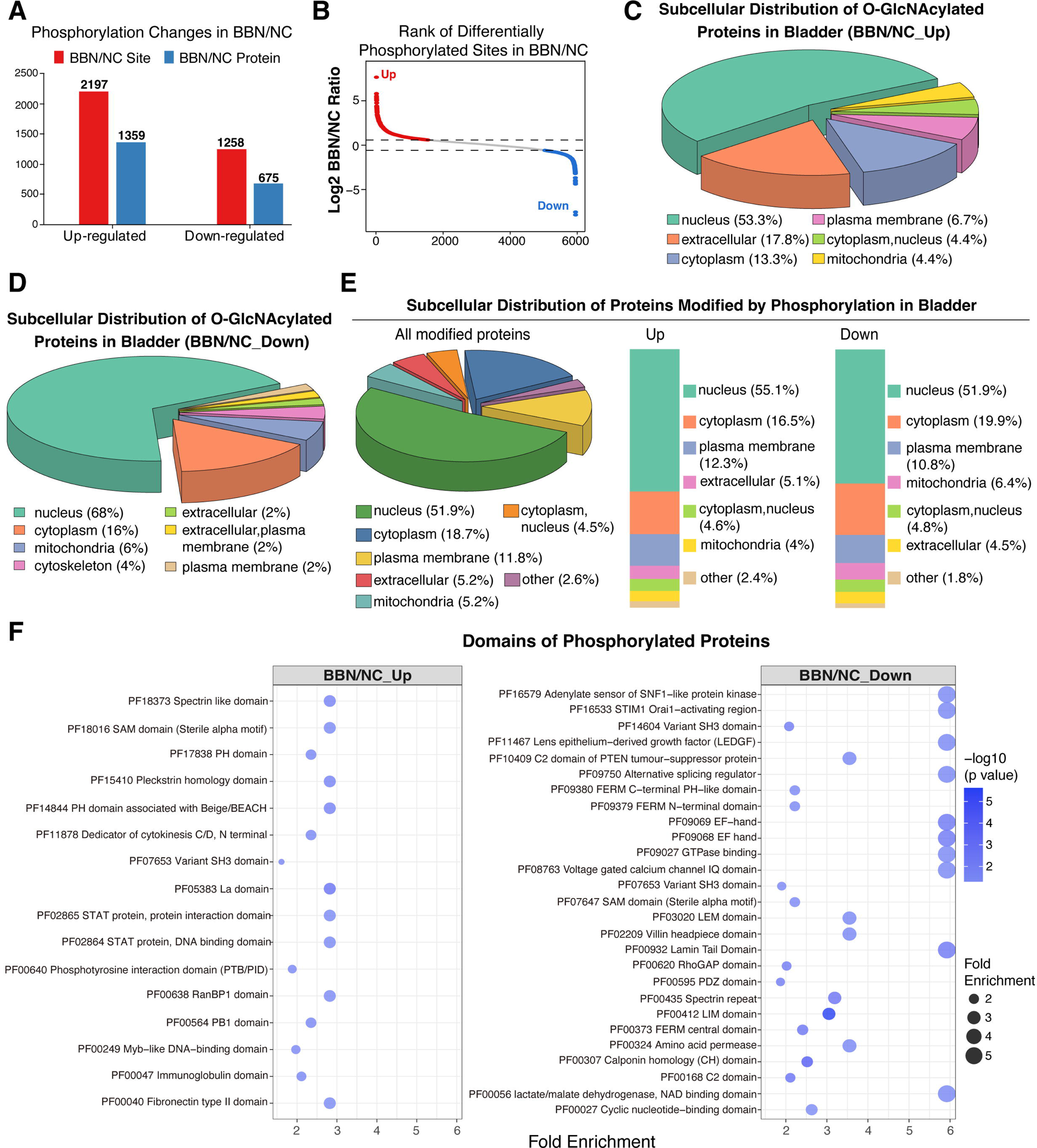

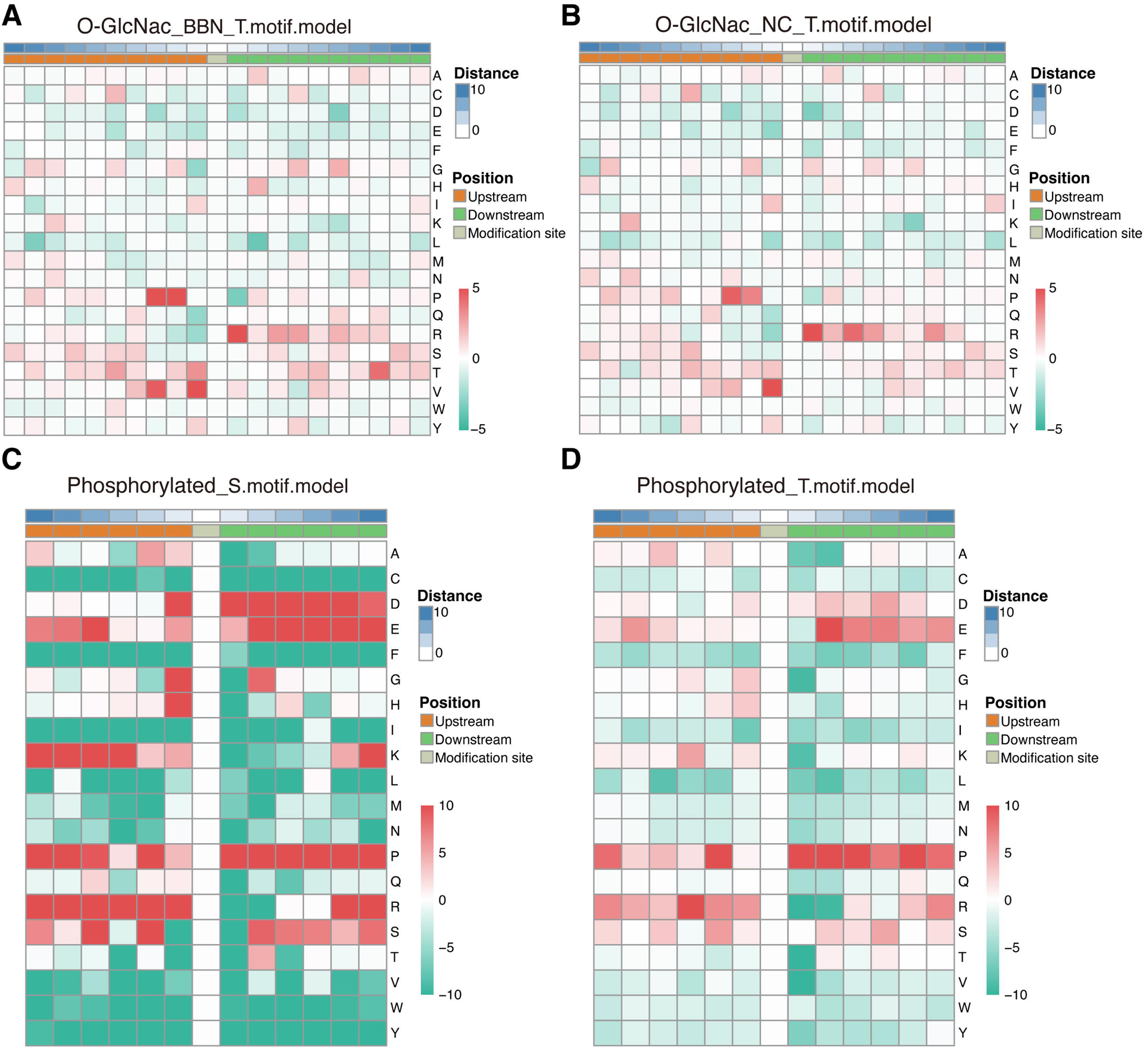

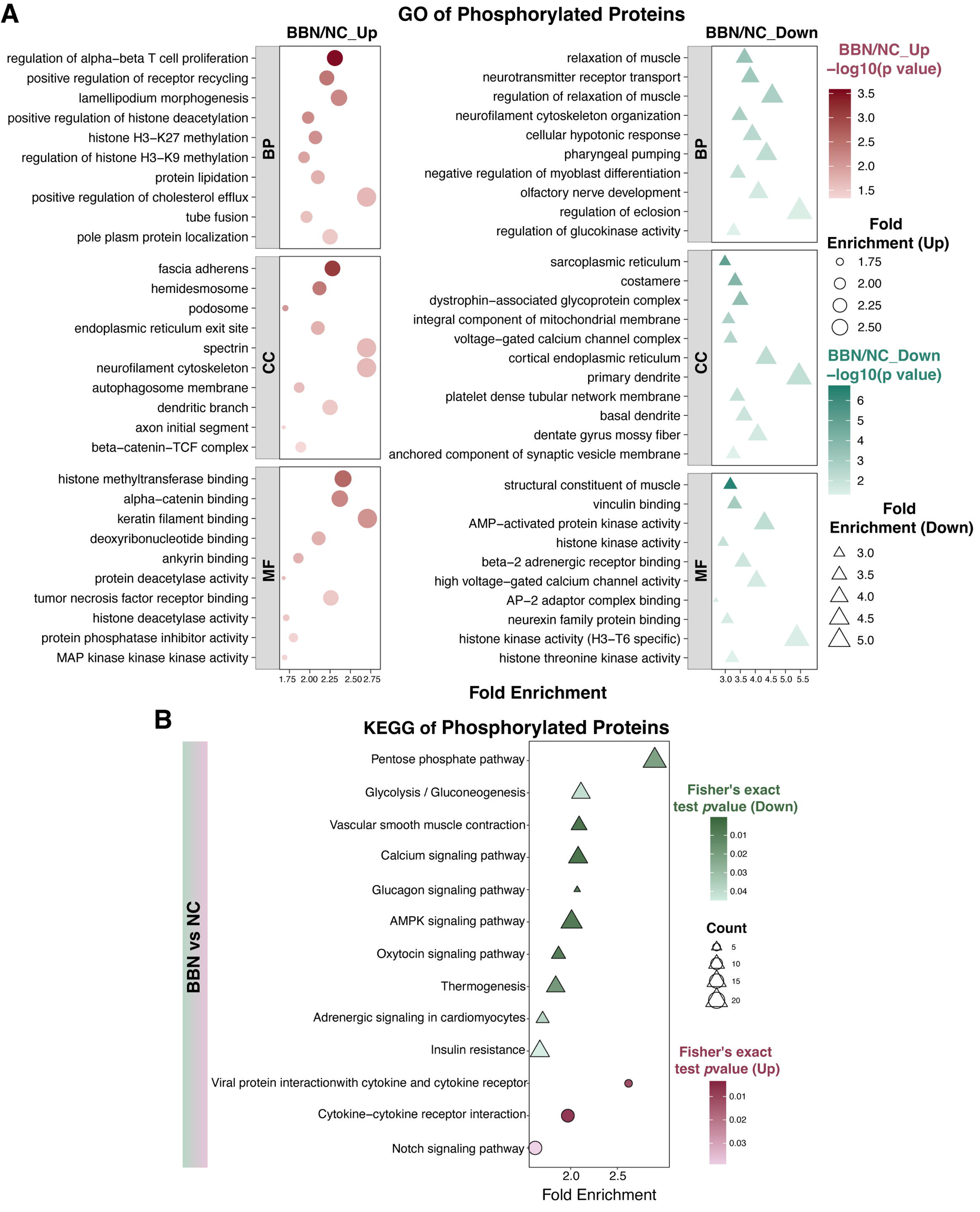

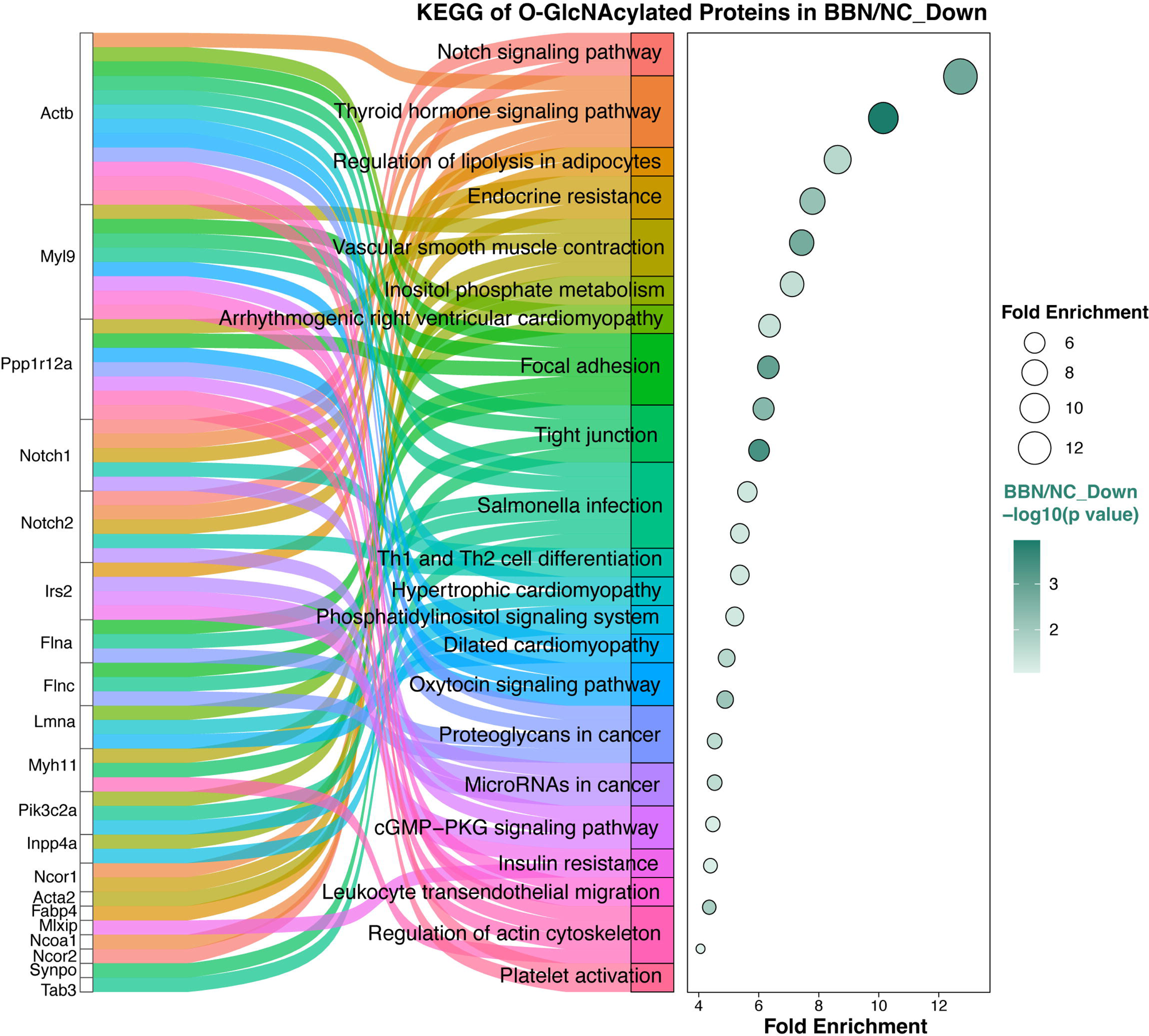

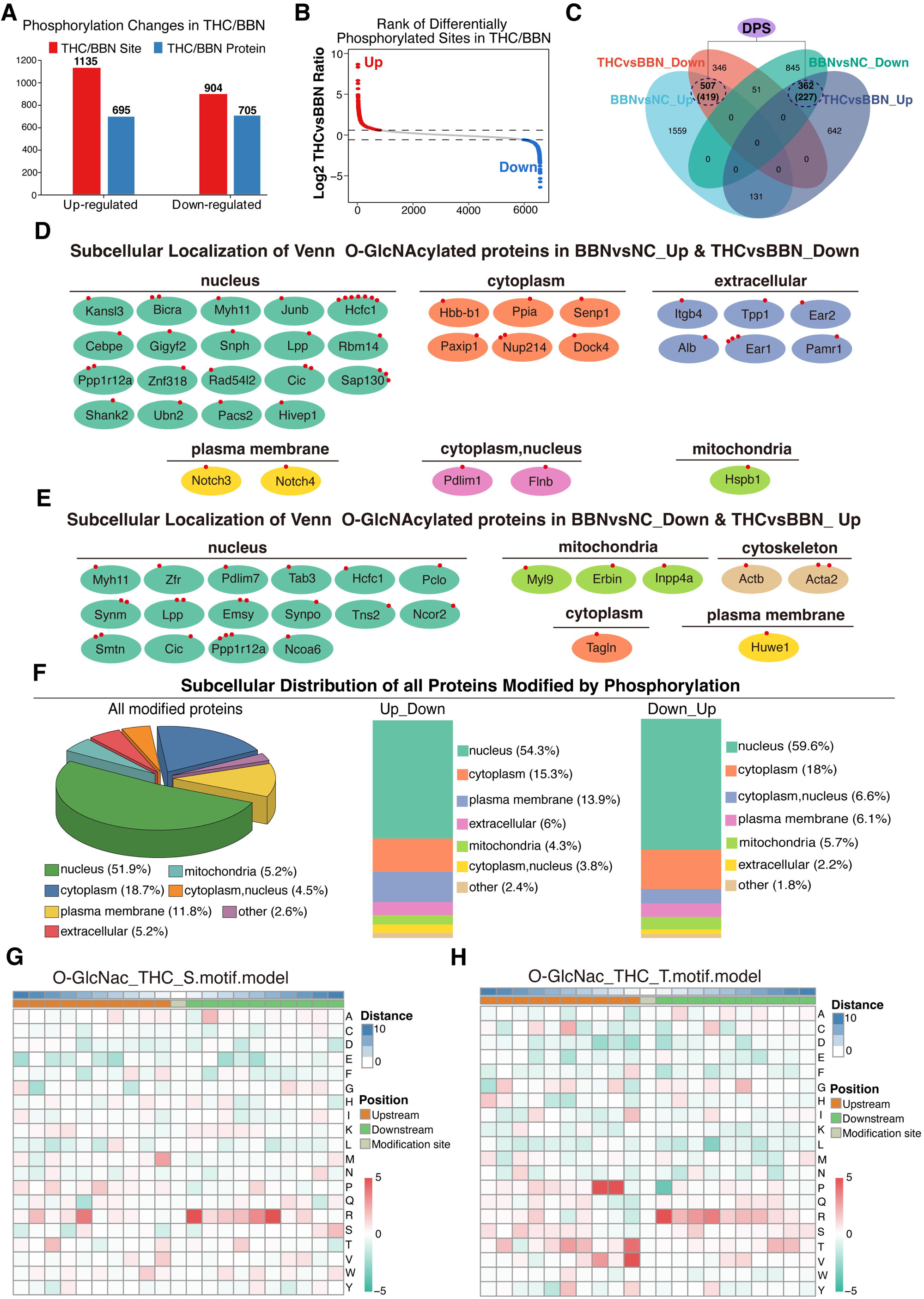

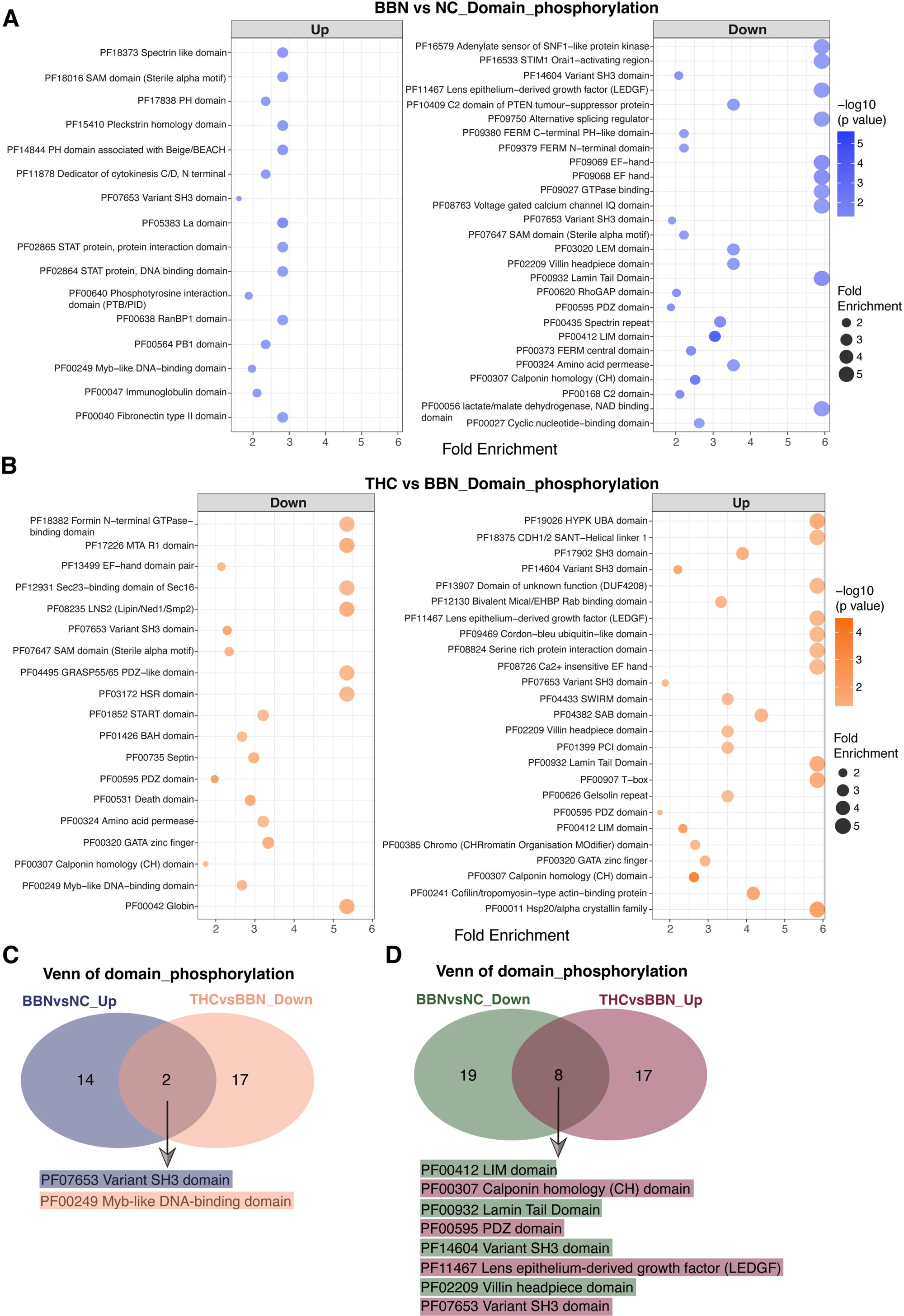

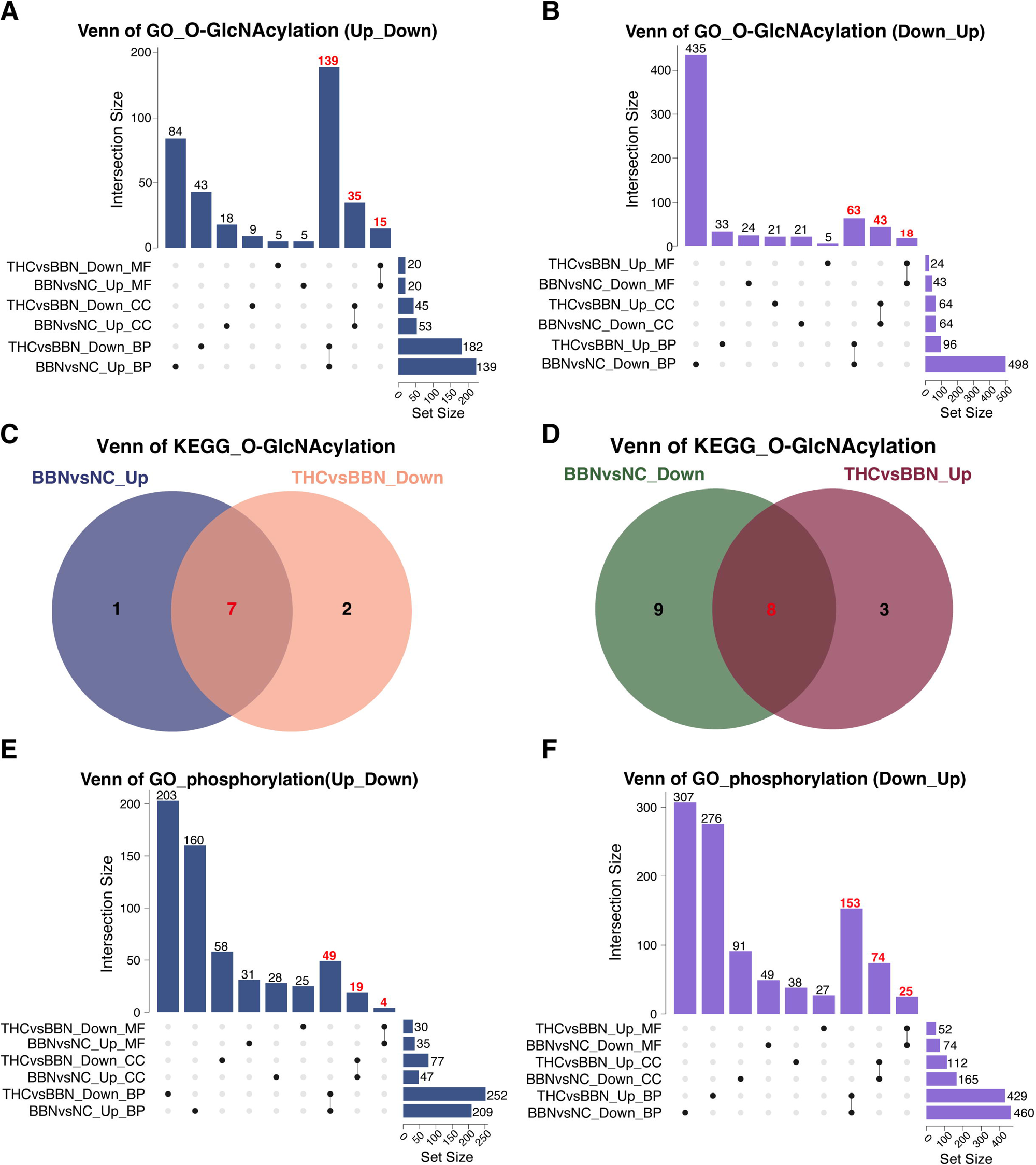

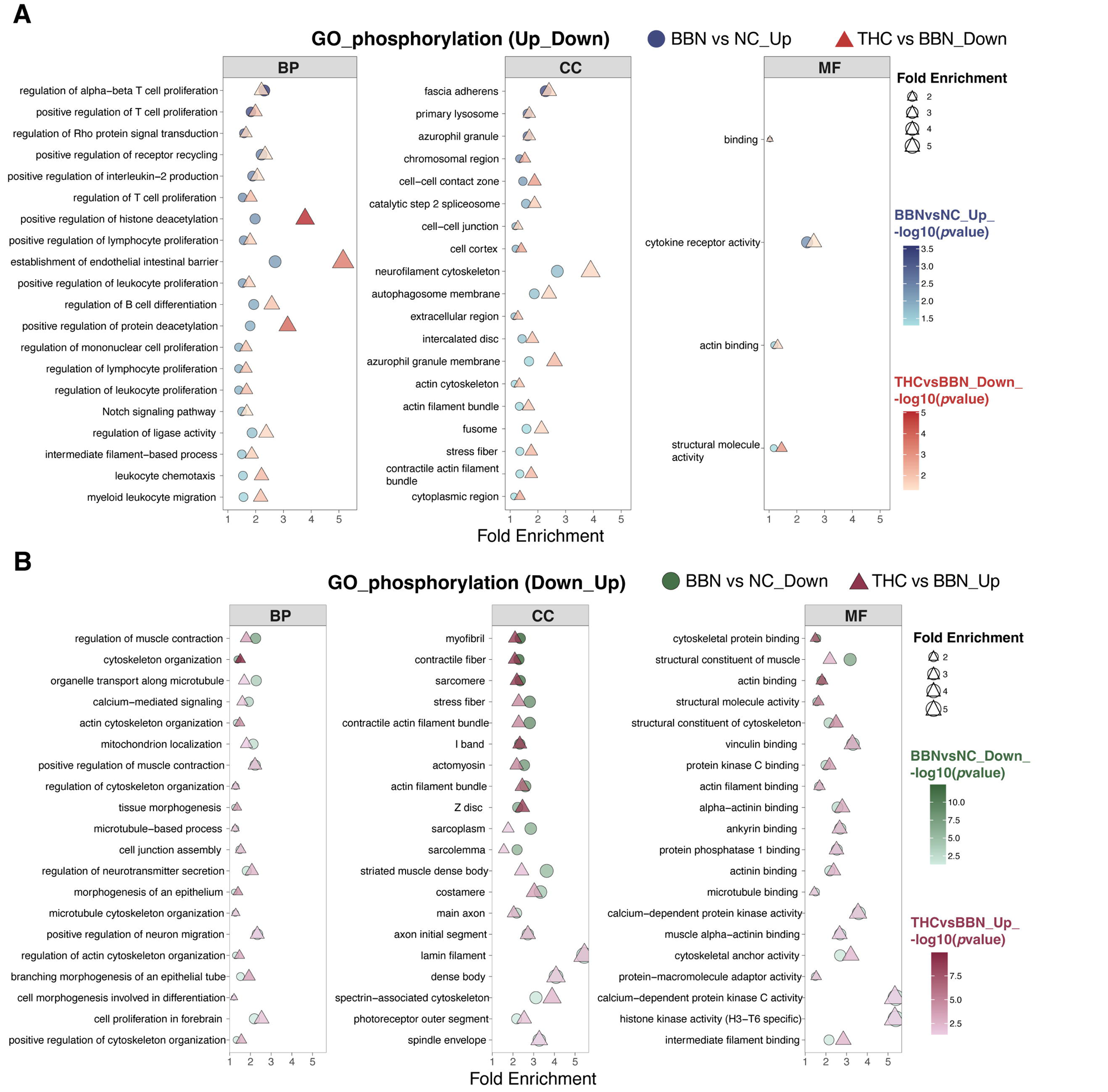

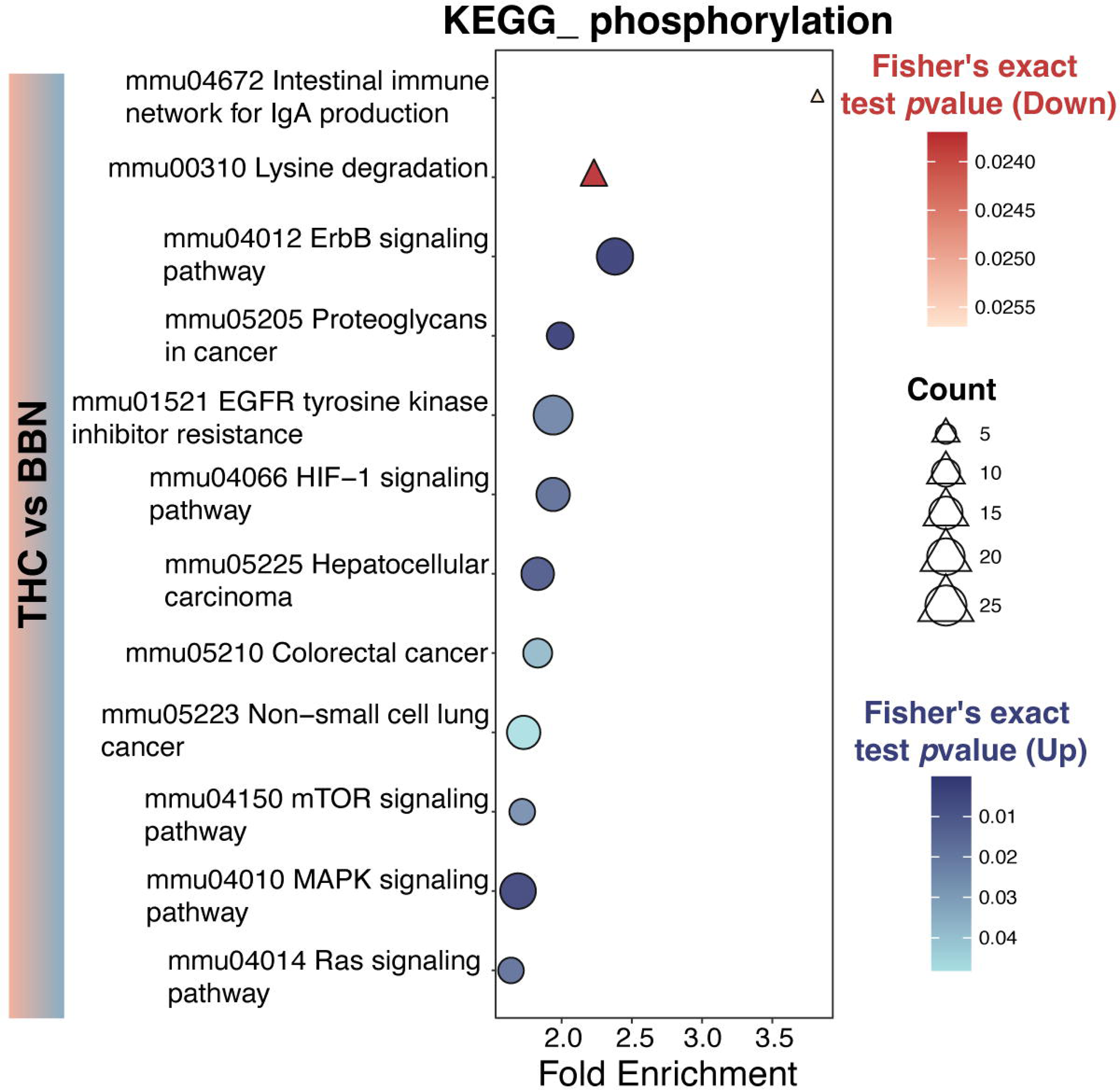

## Notes

### Competing Interest Statement

The authors have declared no competing interest.

