## Supplementary Information for "Tetrahydrocurcumin Suppresses Bladder Carcinogenesis via Reprogramming O-GlcNAcylation-Phosphorylation Crosstalk"

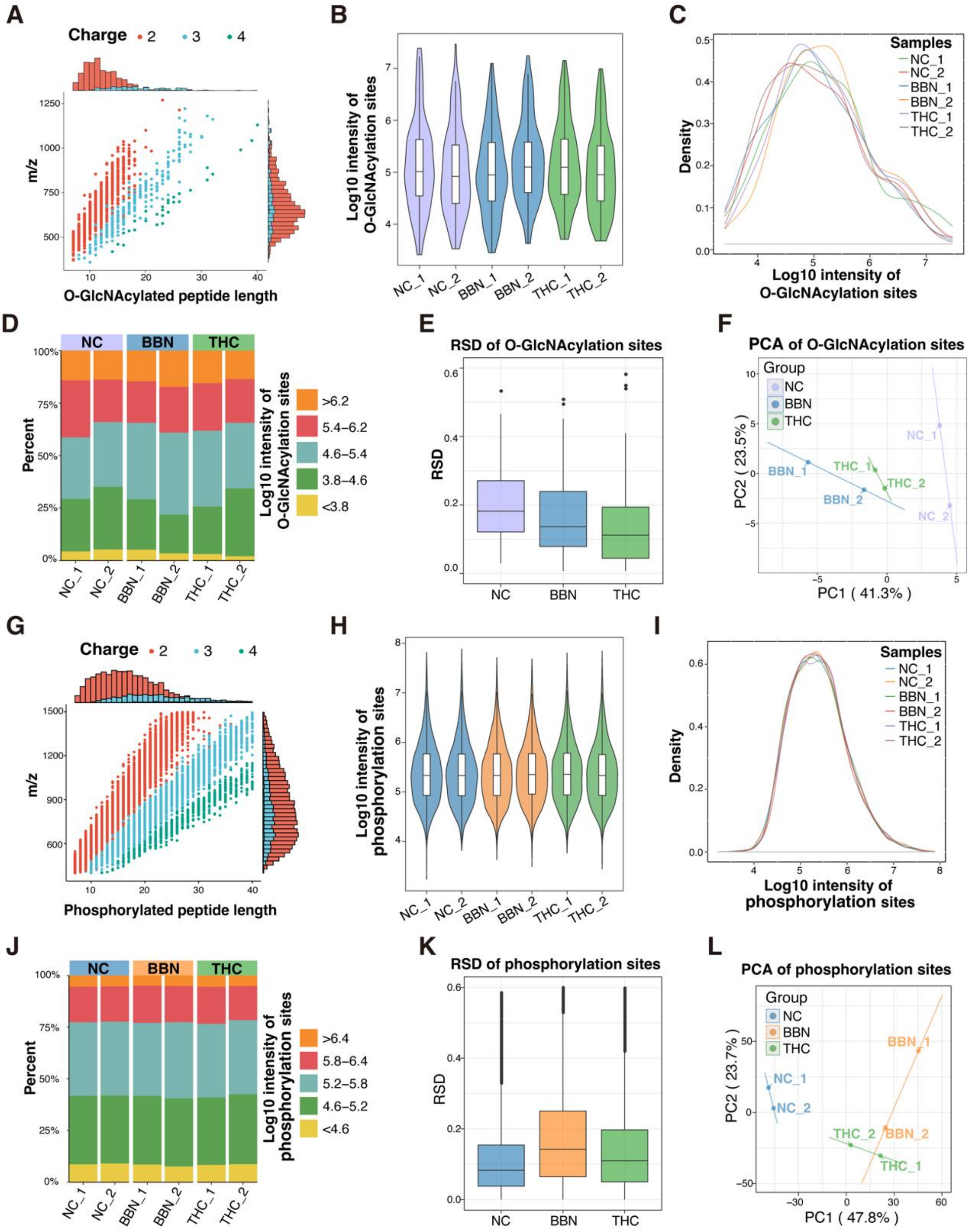

Figure S1 Quality control of protein O-GlcNAcylation or phosphorylation. **(A)** Distribution of m/z values and peptide lengths of identified O-GlcNAcylated peptides. Different colors indicate charge states (2+, 3+, and 4+). Marginal histograms show the overall distribution of peptide length and m/z. **(B)** Violin plots showing the log10 intensity distribution of identified O-GlcNAcylation sites across NC, BBN, and THC groups. **(C)** Density curves of log10 intensities of O-GlcNAcylation sites for each biological replicate. **(D)** Percentage distribution of

O-GlcNAcylation sites stratified by log10 intensity intervals in each sample. **(E)** Relative standard deviation (RSD) of quantified O-GlcNAcylation sites within each group. **(F)** Principal component analysis (PCA) of O-GlcNAcylation site intensities showing sample clustering among NC, BBN, and THC groups. **(G)** Distribution of m/z values and peptide lengths of identified phosphorylated peptides. Different colors indicate charge states (2+, 3+, and 4+). Marginal histograms show the overall distribution of peptide length and m/z. **(H)** Violin plots showing the log10 intensity distribution of identified phosphorylation sites across NC, BBN, and THC groups. **(I)** Density curves of log10 intensities of phosphorylation sites for each biological replicate. **(J)** Percentage distribution of phosphorylation sites stratified by log10 intensity intervals in each sample. **(K)** Relative standard deviation (RSD) of quantified phosphorylation sites within each group. **(L)** Principal component analysis (PCA) of phosphorylation site intensities showing sample clustering among NC, BBN, and THC groups.

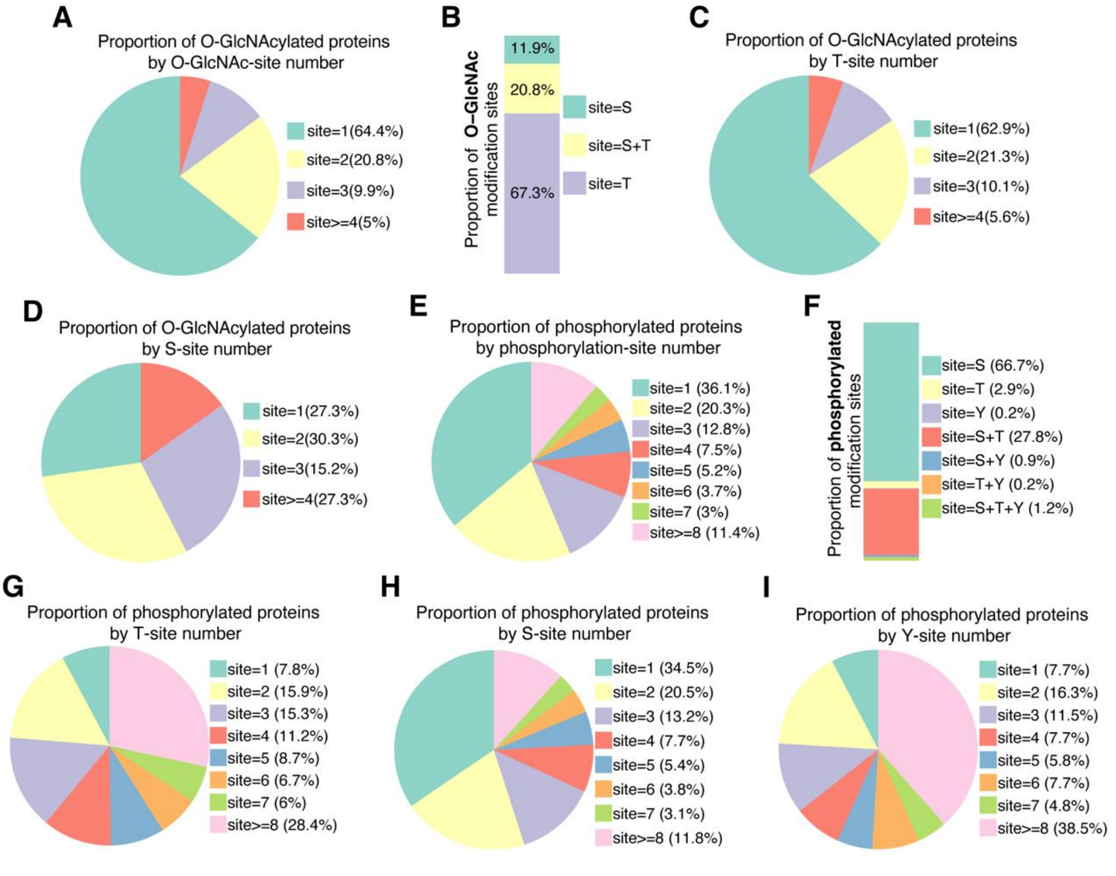


Figure S2. Site-number distribution and residue composition of O-GlcNAcylated and phosphorylated proteins. **(A)** Proportion of O-GlcNAcylated proteins stratified by O-GlcNAc-site number per protein. **(B)** Residue distribution of O-GlcNAc sites, showing the proportions of sites occurring on S (Ser), T (Thr), or both S+T (Ser+Thr). **(C)** Proportion of O-GlcNAcylated proteins by T (Thr)-site number. **(D)** Proportion of O-GlcNAcylated proteins by S (Ser)-site number. (E) Proportion of phosphorylated proteins stratified by phosphorylation-site number per protein. (F) Residue distribution of phosphorylation sites, including S (Ser), T (Thr), and Y (Tyr), and combined categories. **(G)** Proportion of phosphorylated proteins by T (Thr)-site number. **(H)** Proportion of phosphorylated proteins by S (Ser)-site number. **(I)** Proportion of phosphorylated proteins by Y (Tyr)-site number.

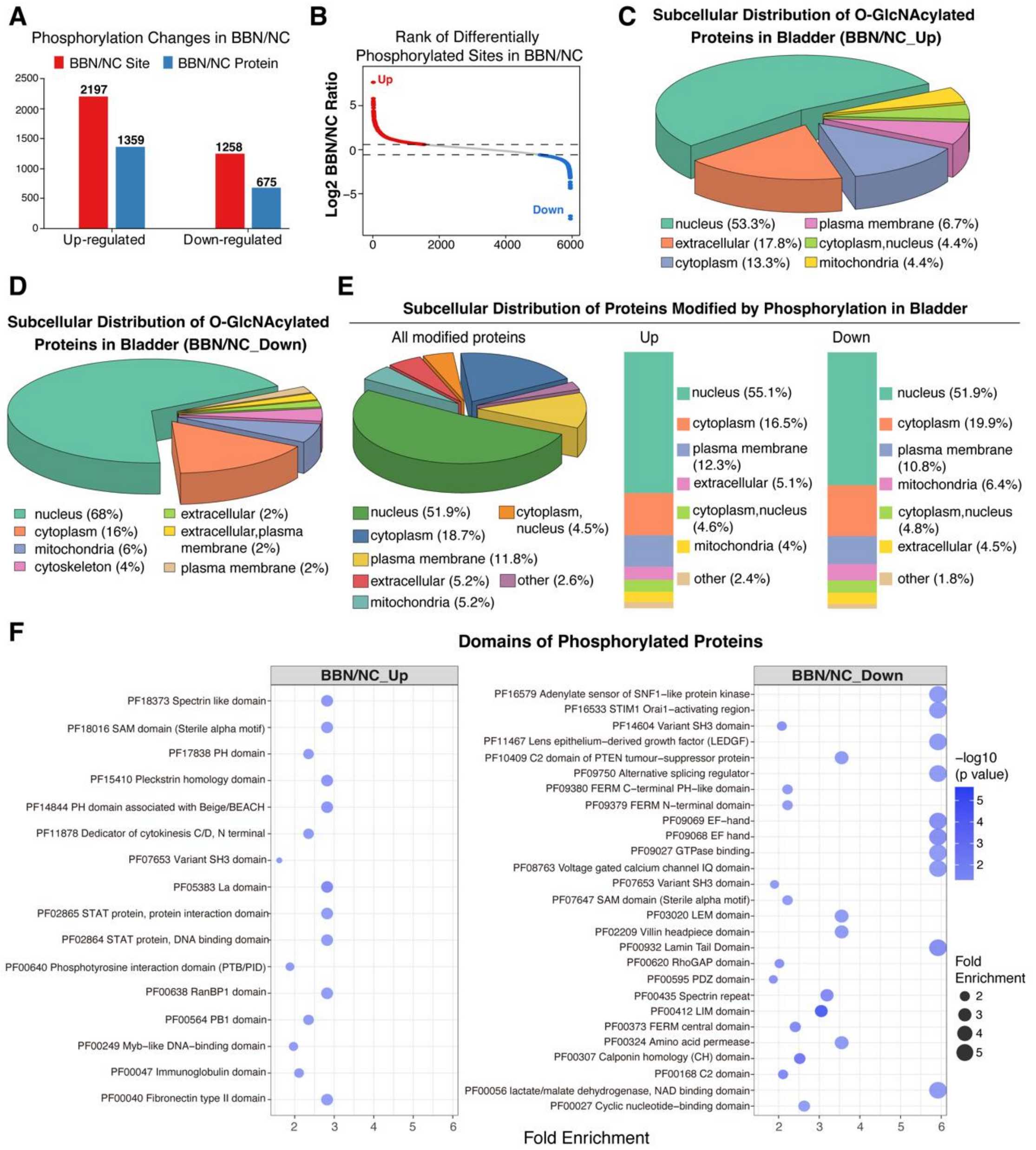

Figure S3. Global characterization of phosphorylation changes in BBN-induced bladder cancer. **(A)** Summary of differentially phosphorylated sites and proteins in the BBN *vs* NC comparisons. **(B)** Rank plots of differential phosphorylated sites in BBN *vs* NC. **(C-D)** Subcellular localization analysis of O-GlcNAcylated proteins that were significantly upregulated (C) or downregulated (D) in BBN *vs* NC. **(E)** Subcellular distribution of phosphorylated proteins in bladder tissues, including all modified proteins, as well as those significantly upregulated or downregulated in BBN *vs* NC. **(F)** Domain enrichment analysis of differentially phosphorylated proteins in BBN/NC_Up and BBN/NC_Down groups.

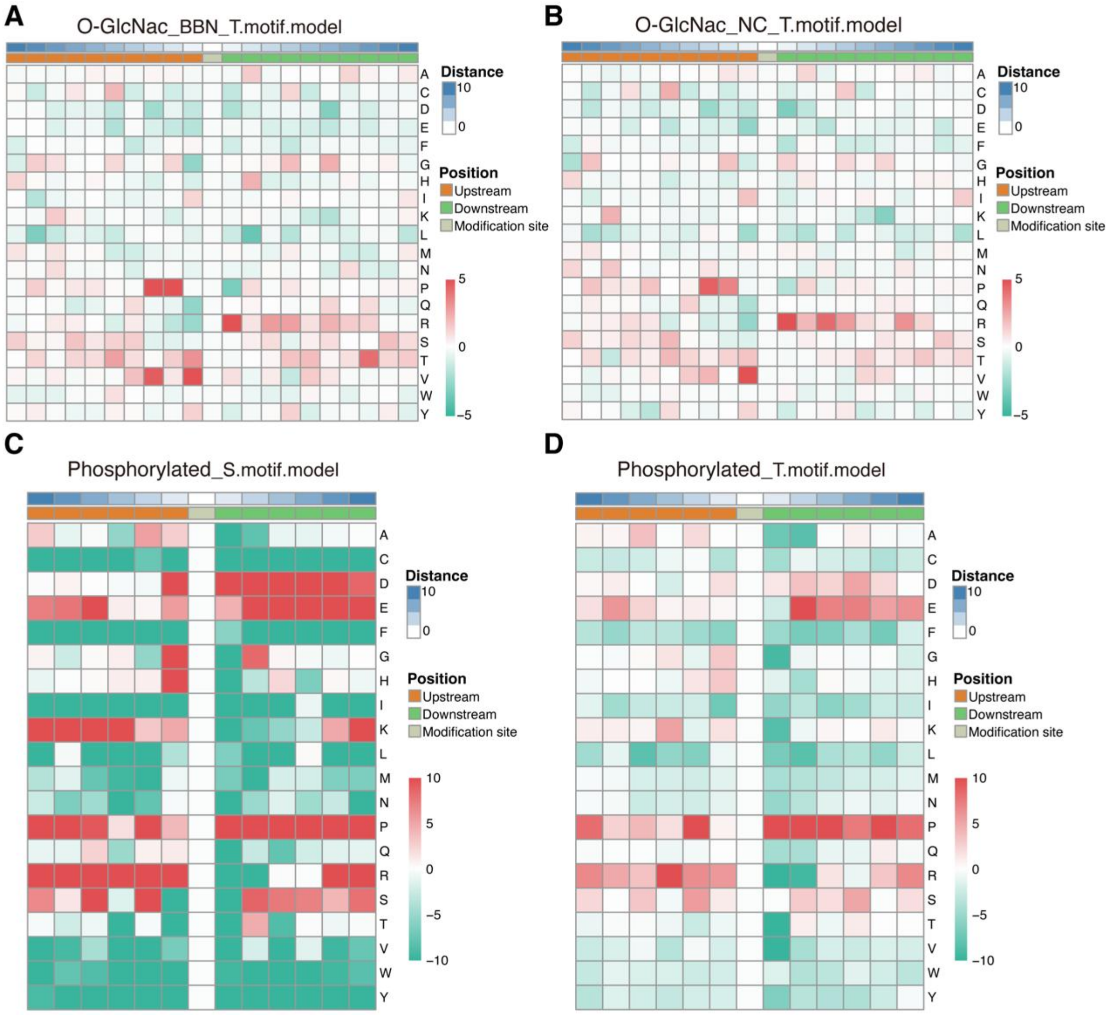


Figure S4 Motif model heatmaps for O-GlcNAcylation and phosphorylation sites. Motif model heatmaps illustrating position-specific residue enrichment/depletion around modification sites. Columns represent positions relative to the modified residue (position 0), and rows represent amino acid types. Color indicates enrichment or depletion relative to background. Heatmaps are shown for O-GlcNAcylation sites (A-B) across NC, BBN and for phosphorylation sites (C-D) as indicated.

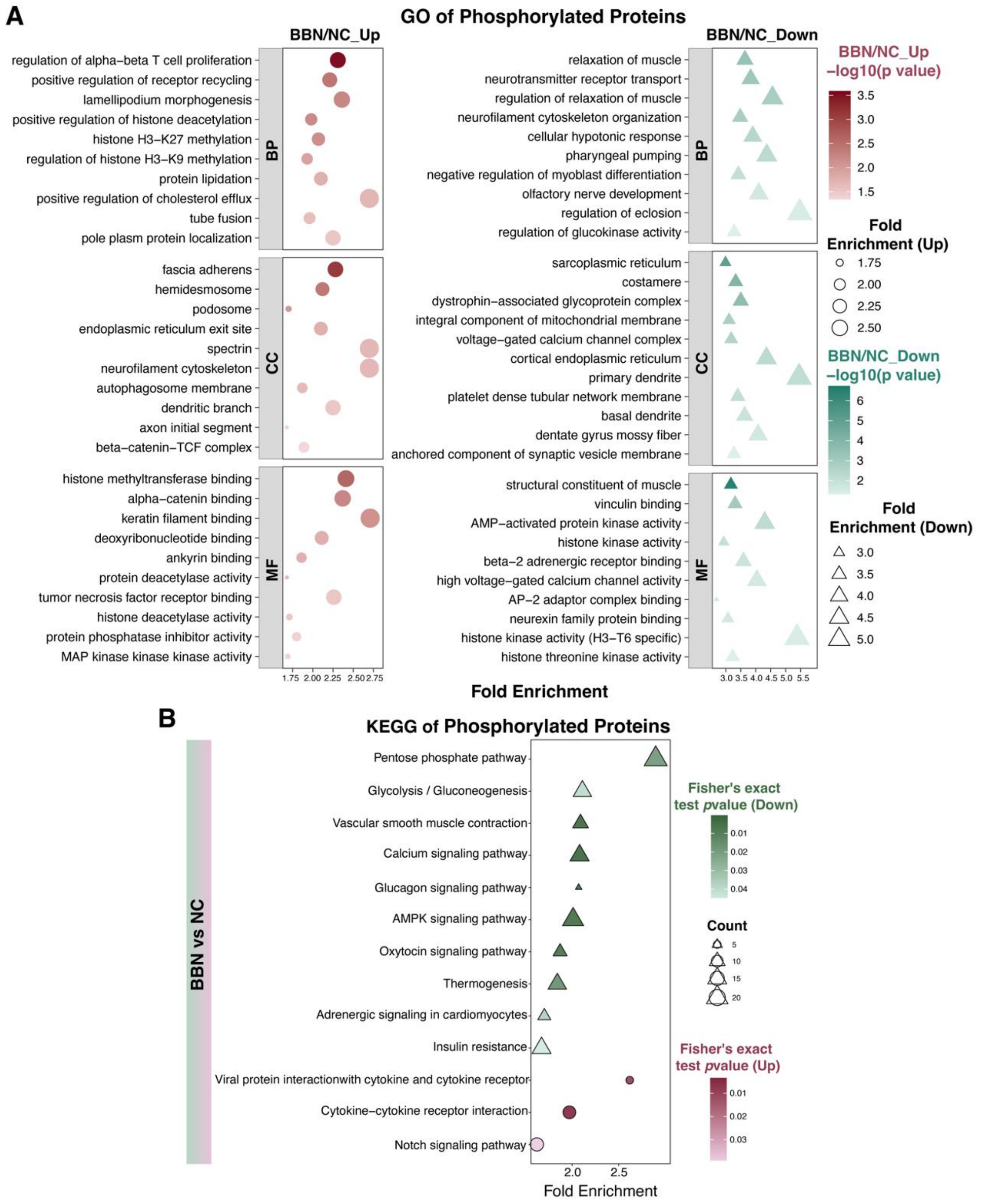


Figure S5 Functional enrichment analysis of differentially phosphorylated proteins in BBN *vs* NC bladder tissues. **(A)** Gene Ontology (GO) enrichment analysis of phosphorylated proteins that were significantly upregulated (BBN/NC_Up) or downregulated (BBN/NC_Down) in the BBN group compared with NC. Biological Process (BP), Cellular Component (CC), and Molecular

Function (MF) categories are shown. **(B)** KEGG pathway enrichment analysis of phosphorylated proteins in the BBN/NC group.

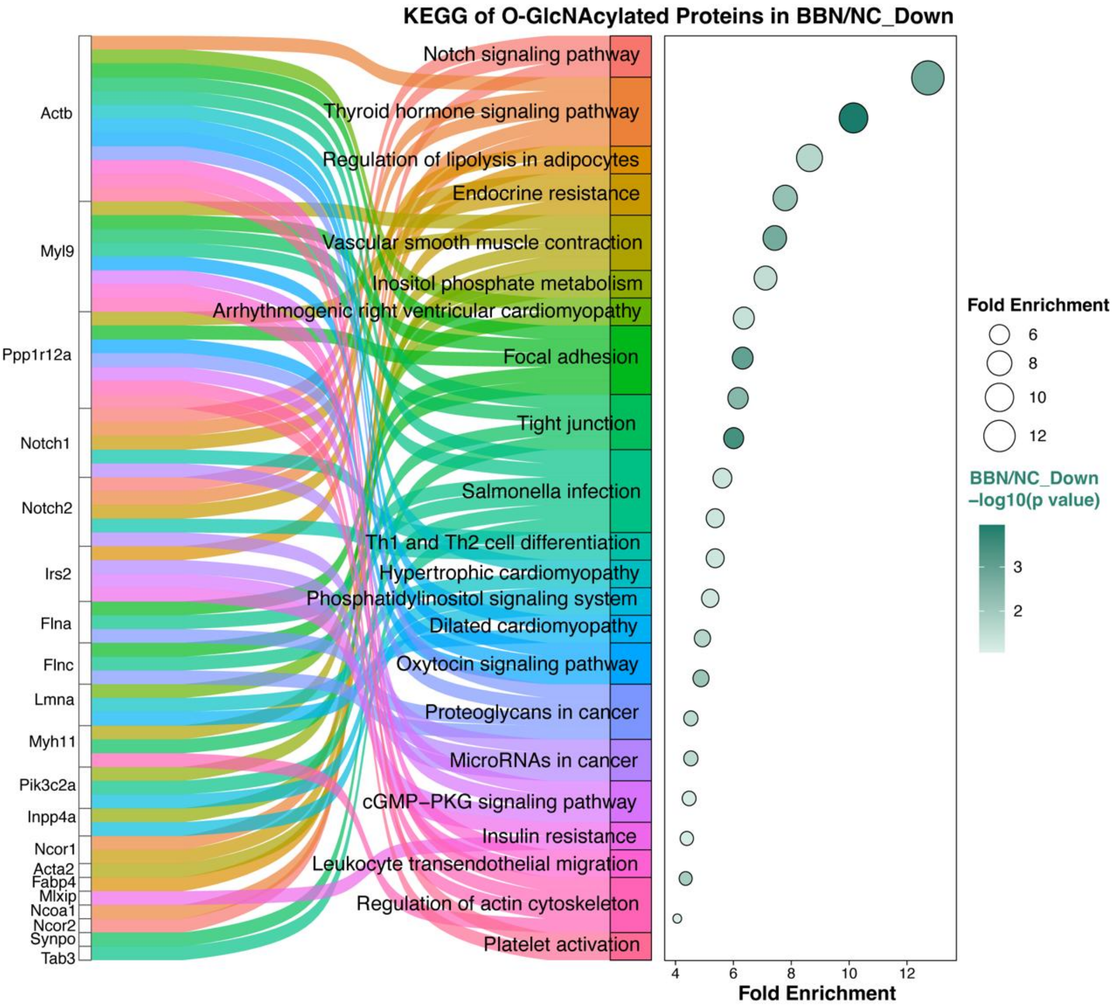


Figure S6 KEGG pathway enrichment analysis of O-GlcNAcylated proteins in the BBN/NC_Down group.

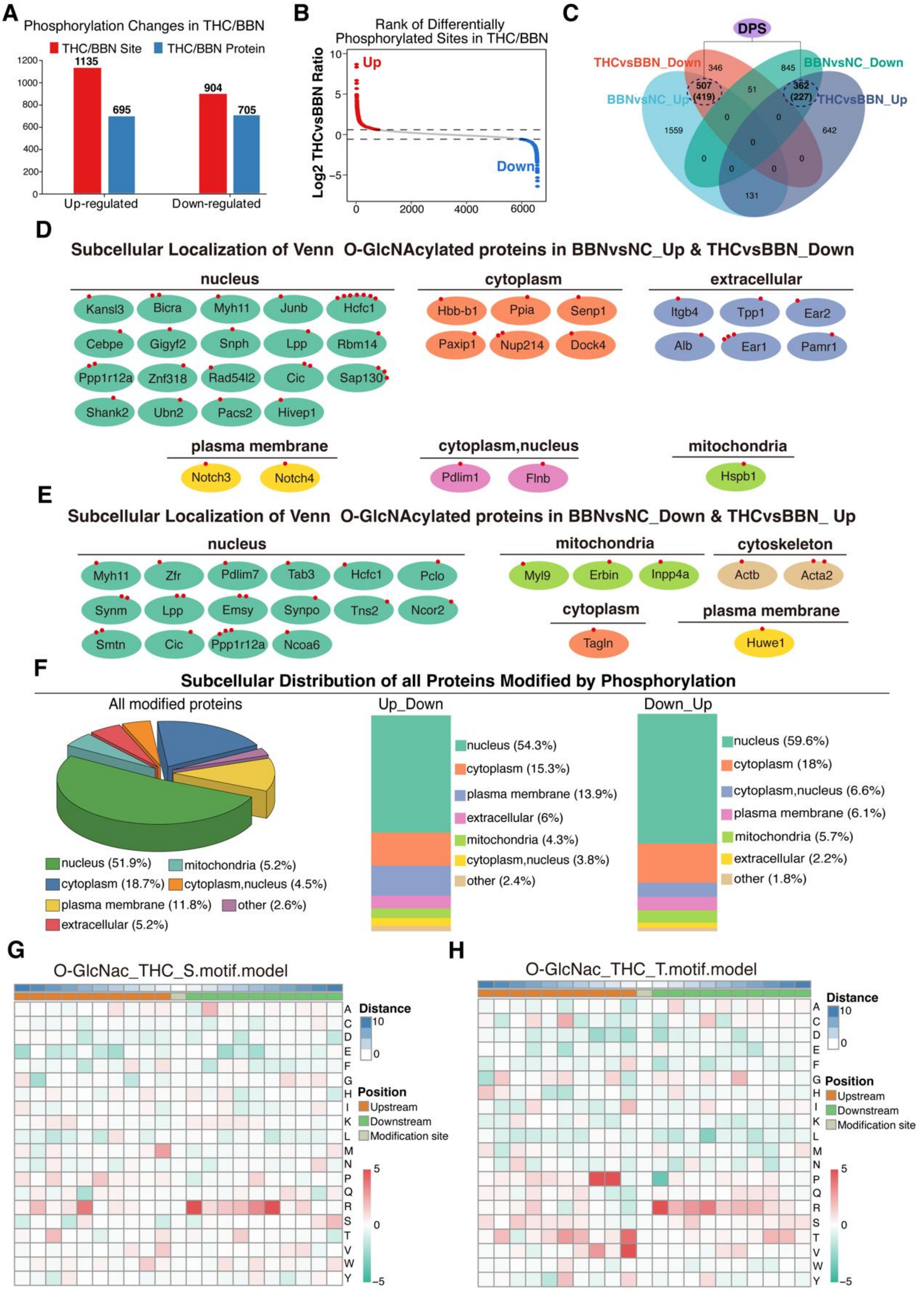


Figure S7. **(A)** Summary of differentially phosphorylated sites and proteins in the THC *vs* BBN comparison. **(B)** Rank plots of differential phosphorylated sites in THC *vs* BBN comparison. **(C)** Intersection analysis of directionally regulated differentially phosphorylated sites (DPS) across BBN vs NC and THC vs BBN, stratified by up/down regulation patterns; numbers indicate sites (proteins). **(D)** Subcellular localization of overlapping O-GlcNAcylated proteins with the BBN *vs* NC_Up and THC *vs* BBN_Down pattern. **(E)** Subcellular localization of overlapping O-GlcNAcylated proteins with the BBN *vs* NC_Down and THC *vs* BBN _Up pattern. **(F)** Subcellular localization distribution of all identified phosphorylated proteins. **(G-H)** Motif model analysis of sequence environments surrounding O-GlcNAcylated residues following THC treatment. Heatmap showing amino acid enrichment patterns around O-GlcNAcylated serine (S) residues identified in the THC dataset (G). Heatmap showing amino acid enrichment patterns around O-GlcNAcylated threonine (T) residues (H).

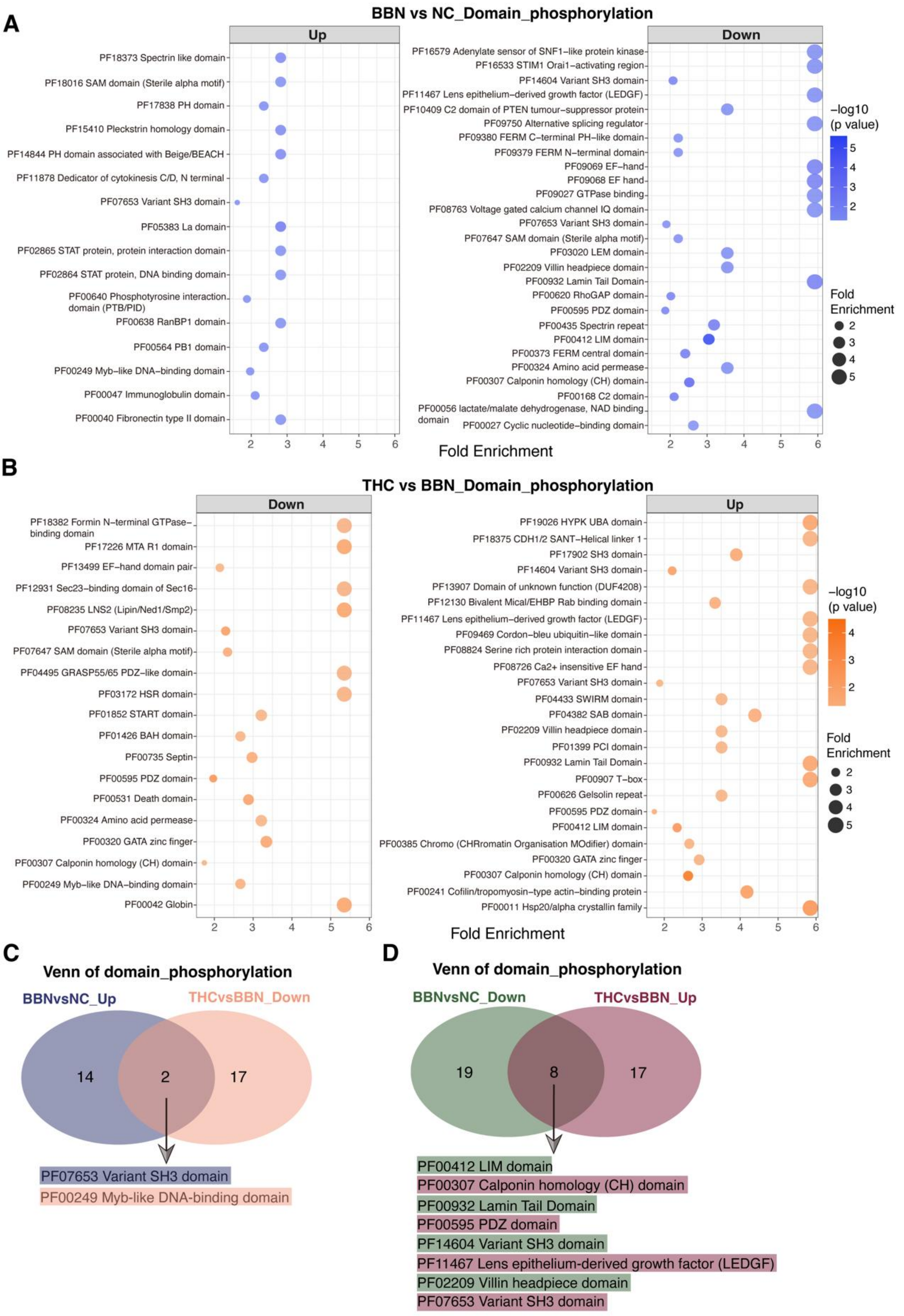


Figure S8. Protein domain enrichment analysis of differentially phosphorylated proteins. **(A)** Bubble plot showing protein domain enrichment for proteins with increased or decreased phosphorylation in the BBN vs NC comparison. **(B)** Bubble plot showing protein domain enrichment for proteins with increased or decreased phosphorylation in the THC vs BBN comparison. **(C)** Venn diagram showing the overlap of enriched domains between the BBN vs NC Up and THC vs BBN Down comparisons. **(D)** Venn diagram showing the overlap of enriched domains between the BBN vs NC Down and THC vs BBN Up comparisons.

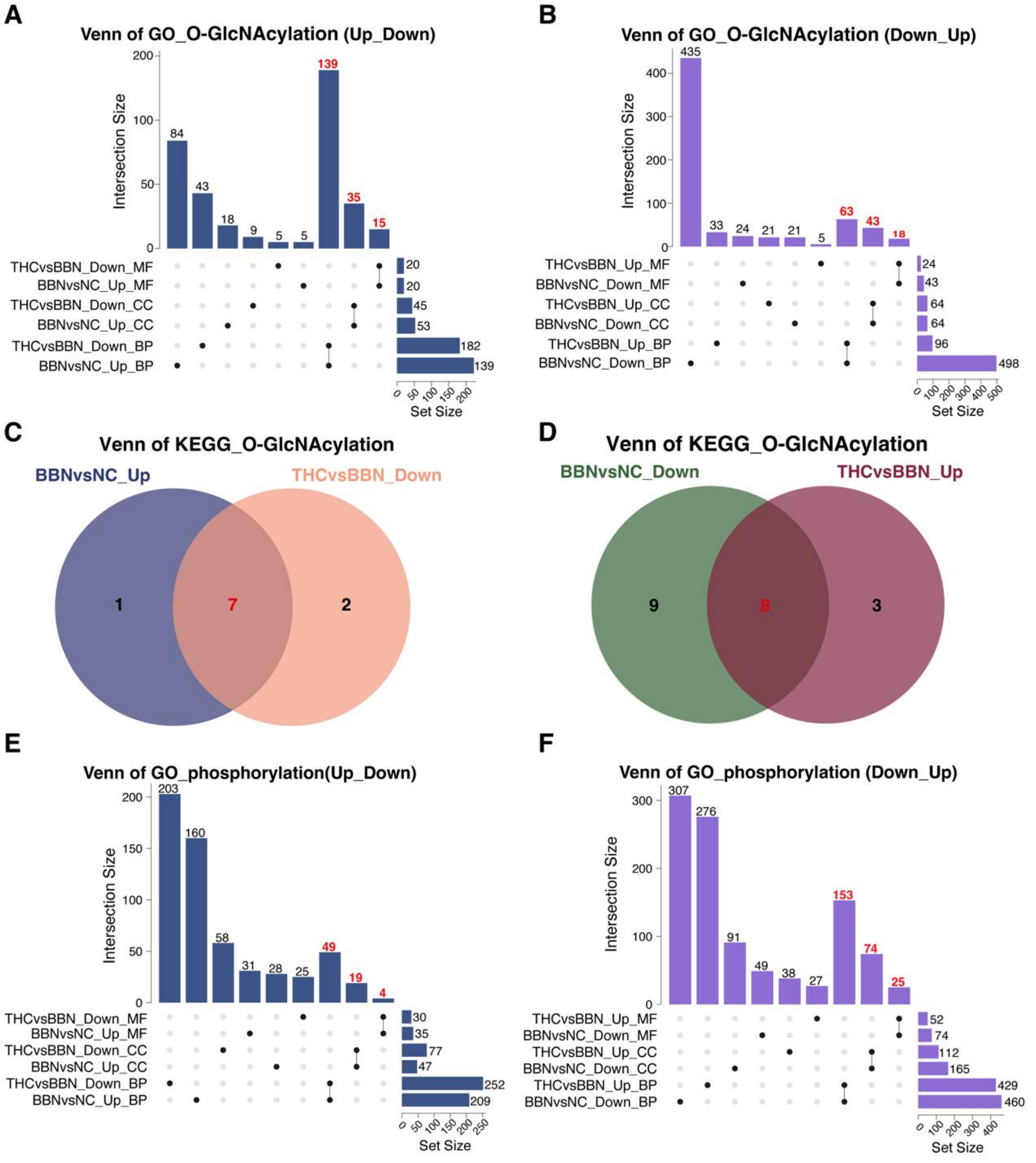


Figure S9. Intersection analysis of GO terms and KEGG pathways for O-GlcNAcylation- and phosphorylation-associated proteins. **(A-B)** UpSet plots showing overlaps of enriched GO terms for differentially O-GlcNAcylated proteins in the Up_Down (BBN vs NC_up / THC vs BBN_down) (A) and Down_Up (BBN vs NC_down / THC vs BBN_up) (B) patterns across BP,

CC, and MF categories. **(C-D)** Venn diagrams showing overlaps of enriched KEGG pathways for O-GlcNAcylation in the Up_Down (C) and Down_Up (D) patterns. **(E-F)** UpSet plots showing overlaps of enriched GO terms for differentially phosphorylated proteins in the Up_Down (E) and Down_Up (F) patterns.

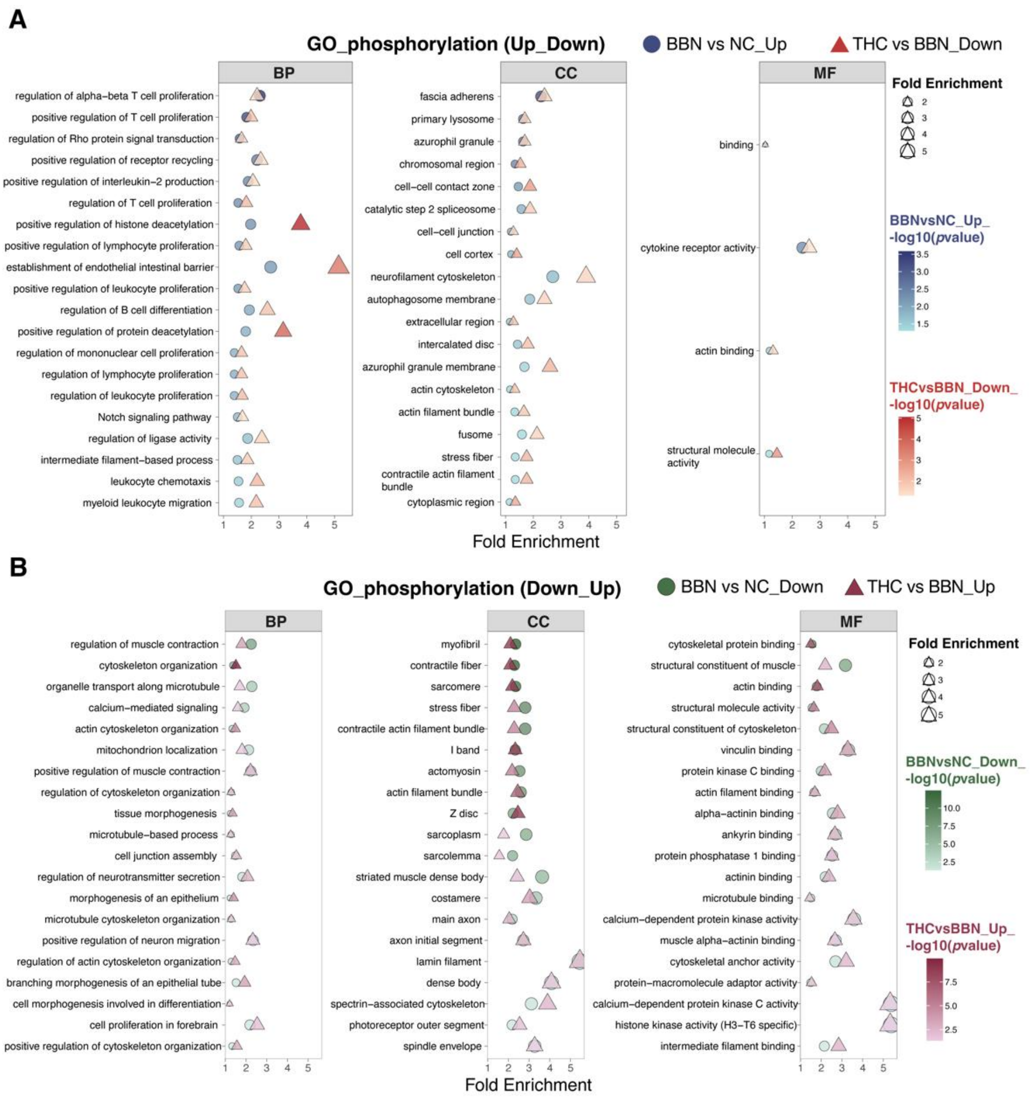


Figure S10. GO enrichment analysis of differentially phosphorylated proteins. **(A)** GO enrichment results for the Up_Down pattern (BBN vs NC_up and THC vs BBN_down) across BP, CC, and MF categories. **(B)** GO enrichment results for the Down_Up pattern (BBN vs NC_down and THC vs BBN_up) across BP, CC, and MF categories.

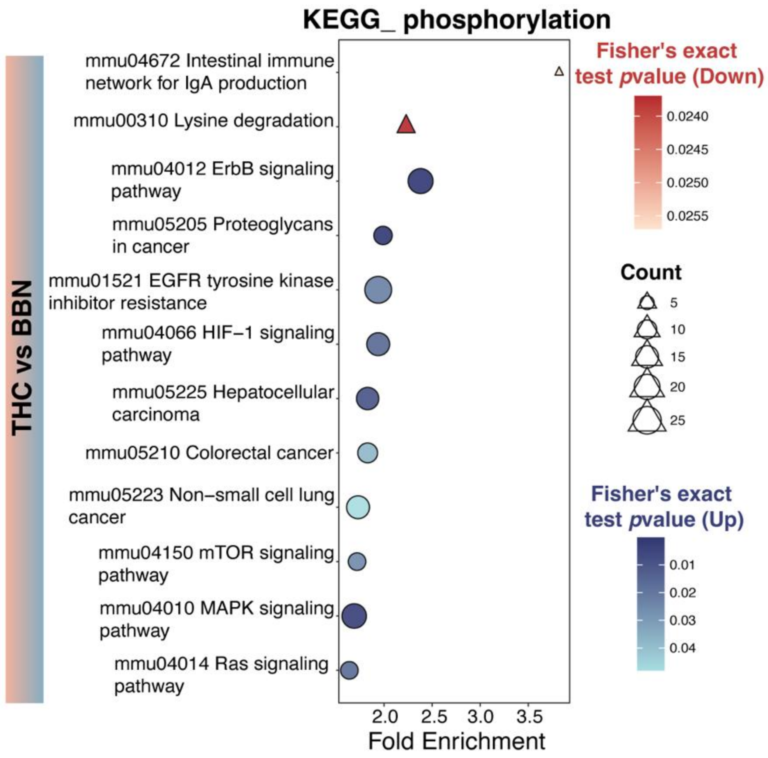


Figure S11. KEGG enrichment results for differentially phosphorylated proteins in THC vs BBN.
